# *doubletrouble:* an R/Bioconductor package for the identification, classification, and analysis of gene and genome duplications

**DOI:** 10.1101/2024.02.27.582236

**Authors:** Fabricio Almeida-Silva, Yves Van de Peer

## Abstract

Gene and genome duplications are major evolutionary forces that shape the diversity and complexity of life. However, different duplication modes have distinct impacts on gene function, expression, and regulation. Existing tools for identifying and classifying duplicated genes are either outdated or not user-friendly. Here, we present *doubletrouble*, an R/Bioconductor package that provides a comprehensive and robust framework for analyzing duplicated genes from genomic data. *doubletrouble* can detect and classify gene pairs as derived from six duplication modes (segmental, tandem, proximal, retrotransposon-derived, DNA transposon-derived, and dispersed duplications), calculate substitution rates, detect signatures of putative whole-genome duplication events, and visualize results as publication-ready figures. We applied *doubletrouble* to classify the duplicated gene repertoire in 822 eukaryotic genomes, which we made available through a user-friendly web interface (available at https://almeidasilvaf.github.io/doubletroubledb). *doubletrouble* is freely accessible from Bioconductor (https://bioconductor.org/packages/doubletrouble), and it provides a valuable resource to study the evolutionary consequences of gene and genome duplications.

## Introduction

Gene and genome duplications are an important source of novel genetic material for evolution to work with (Ohno 1970). However, gene and genome duplications contribute to genome evolution in different ways, and genes created by these two mechanisms evolve under different selection pressures, and display different evolutionary trajectories at the genomic, transcriptomic, and epigenomic levels (Casneuf et al. 2006; Freeling and Thomas 2006; Ganko et al. 2007; Hakes et al. 2007; Freeling 2009; De Smet et al. 2013; Li et al. 2016; Qiao et al. 2018; Qiao et al. 2019; Almeida-Silva et al. 2020; Almeida-Silva and Van de Peer 2023; Kenchanmane Raju et al. 2023; Lallemand et al. 2023). Furthermore, depending on the duplication mode, gene retention can be biased, with some gene classes (e.g., transcription factors and members of protein complexes) being preferentially retained after whole-genome duplication (Freeling and Thomas 2006; Birchler and Veitia 2010).

To understand the differential contribution of each duplication mode to genome evolution and the emergence of novel traits, researchers typically use a utility program in the MCScanX toolkit (Wang et al. 2012) to classify gene pairs as derived from segmental, tandem, proximal, or dispersed duplications. However, MCScanX is no longer actively maintained, and its classification scheme does not include transposon-derived duplications. The *DupGen_finder* pipeline (Qiao et al. 2019) extends the classification scheme in MCScanX to identify transposon-derived duplicates, but it consists of a set of Perl scripts, not a distributable software package (*i*.*e*., available from a package manager, with unit tests, continuous integration, and documentation), hindering its stability and usability.

Here, we introduce *doubletrouble*, an R/Bioconductor package to identify, classify, and analyze duplicated genes from genomic data. *doubletrouble* can identify and classify duplicated gene pairs as derived from segmental, tandem, proximal, retrotransposon-derived, DNA transposon-derived, and dispersed duplications. Duplicate pairs identified and classified with *doubletrouble* can be further used to calculate substitution rates (K_a_, K_s_, and their ratio K_a_/K_s_), investigate signatures of potential whole-genome duplication events, and generate publication-ready figures. We demonstrate *doubletrouble*’s effectiveness by identifying and classifying the entire duplicated gene repertoire in 822 eukaryotic genomes from Ensembl instances. Finally, to facilitate data reuse, we created a web application (https://almeidasilvaf.github.io/doubletroubledb) where users can explore and download data generated here on the duplicated gene repertoire across the Eukarya tree of life.

## Implementation

*doubletrouble* is part of the Bioconductor ecosystem of R packages and, as such, was designed to easily interoperate with other Bioconductor packages. For that, data classes used in *doubletrouble* are either base R classes or core Bioconductor S4 classes, such as *GRanges* objects to store genomic annotation (Lawrence et al. 2013), and *AA*/*DNAStringSet* objects to store sequence data (Pagès et al. 2023).

### Data input and processing

The required input data are whole-genome protein sequences (one per gene, primary transcript only) as *AAStringSet* objects, and gene annotation as *GRanges* objects, both in lists with matching names representing species names or any other genome identifiers. Users can create such lists of *AAStringSet* and *GRanges* objects from multiple FASTA and GFF/GTF files in a directory with the functions *fasta2AAStringSetlist()* and *gff2GRangesList()*, available in the R package syntenet (Almeida-Silva et al. 2023). Sequence and annotation data must be processed with the function *process_input()* from the syntenet package to ensure that only one sequence per gene is present (see Almeida-Silva et al. (2023) for details).

### Identification and classification of duplicated gene pairs

Processed data can be used as input to the function *run_diamond()* from the syntenet package to perform intraspecies similarity searches with DIAMOND (Buchfink et al. 2021) (default parameters: top hits = 5; E-value = 1e-10). Alternatively, users who wish to use a different similarity search program (e.g., BLAST (Altschul et al. 1997) or MMseqs2 (Steinegger and Söding 2017)) can export processed sequences to FASTA files with the function *export_sequences()*, run their favorite program on the command line, and read its tabular output with the function *read_diamond()*. The tabular output of DIAMOND and similar programs is considered to contain all paralogous gene pairs within a genome (i.e., a “paranome”).

The paranomes and gene annotations for each species are used as input to the function *classify_gene_pairs()*, which classifies paralogs by mode of duplication based on four possible classification schemes (Fig. 1A). In the *binary* scheme, paralogs are classified as derived from either segmental duplications (SD) or small-scale duplications (SSD). The *standard* scheme (default) further classifies SSD-derived pairs as originating from tandem (TD), proximal (PD), or dispersed duplications (DD). The *extended* scheme includes transposon-derived duplications (TRD) as an additional duplication mode, and the *full* scheme further classifies TRD-derived genes as originating from either retrotransposons (rTRD) or DNA transposons (dTRD).

**Fig. 1.**
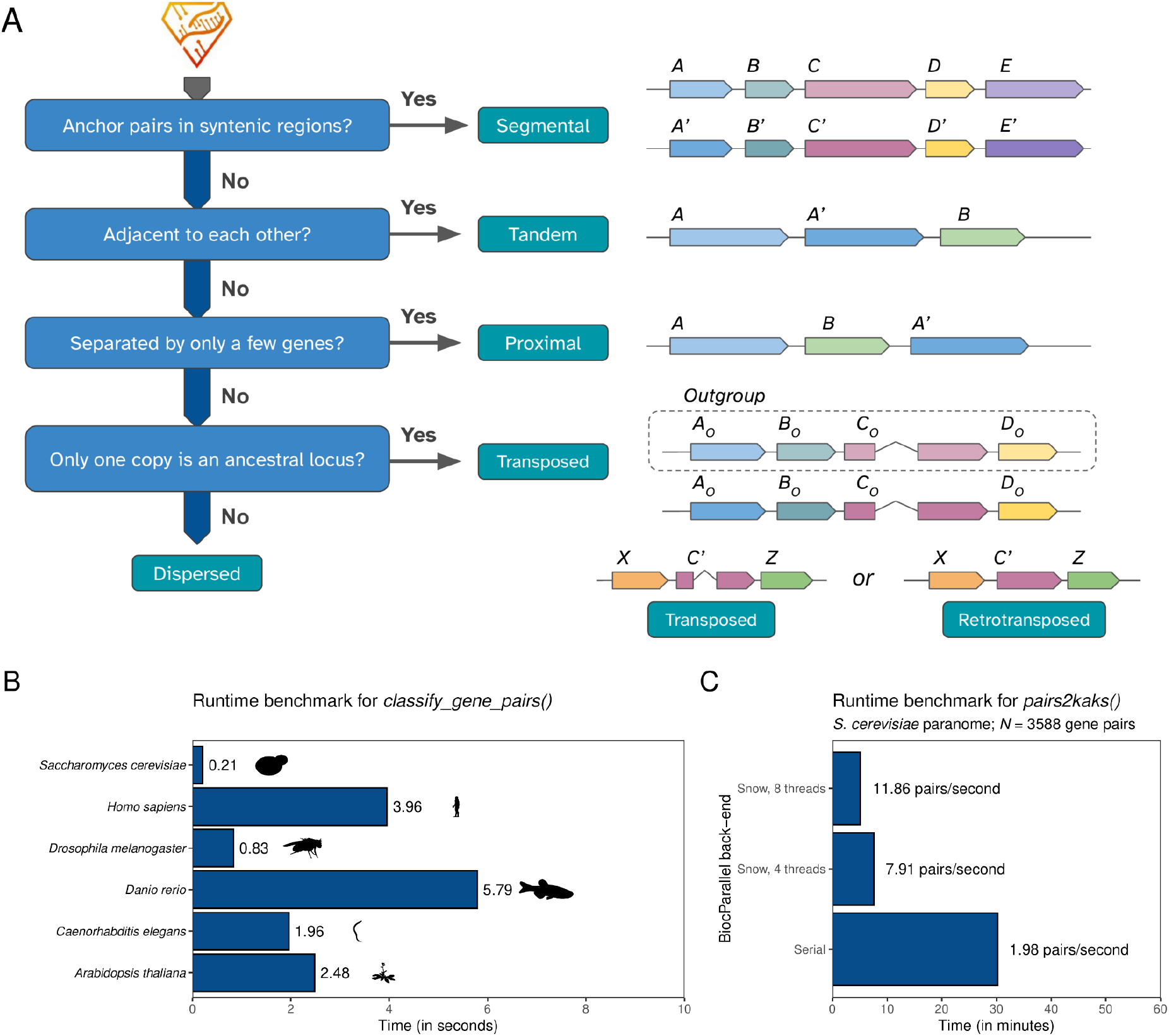
Schematic overview of the classification algorithm for duplicated genes, and runtime benchmarks. **A**. Algorithm for the classification of duplicates. From the output of DIAMOND or related sequence similarity search tools, duplicated gene pairs are classified according to duplication mode in a stepwise manner. Gene pairs that fail to meet the criteria for all duplication modes will be classified as “dispersed duplicates”, which is not a true duplication mechanism, denoting that their duplication mechanism is actually unknown. **B**. Runtime benchmark of the function *classify_gene_pairs()* for model organisms using the ‘standard’ scheme. Silhouettes were obtained from PhyloPic (https://www.phylopic.org/) using the R package rphylopic (Gearty and Jones 2023). **C**. Runtime benchmark of the function *pairs2kaks()* with a single thread and in parallel mode (four and eight threads). For each gene pair, this function performs a pairwise alignment of the sequences, and then calculates substitution rates.

To classify paralogs by duplication mode, the function *classify_gene_pairs()* uses the syntenet package to detect intragenomic syntenic regions. Paralogs that are anchor pairs in syntenic regions are classified as segmental duplicates (SD), and all other duplicates are classified as small-scale duplicates (SSD). SSD pairs that are physically adjacent to each other in the genome are classified as tandem duplicates (TD), SSD pairs that are separated by only a few genes (default = 10, but adjustable) are classified as proximal duplicates (PD), and all other duplicates are classified as dispersed duplicates (DD). To further classify DD pairs as originating from transposon-derived duplications (TRD), users must provide a table with similarity search results between all query species and an outgroup (*e*.*g*., as created with the *run_diamond()* function), which will be used to detect interspecies syntenic regions. DD pairs for which only one copy is an ancestral locus (*i*.*e*., syntenic with the outgroup species) are classified as transposed duplicates (TRD). Finally, TRD pairs for which only one copy is intronless are classified as retrotransposed duplicates (rTRD), while other TRD pairs are classified as DNA transposon-derived duplicates (dTRD).

As genes can be duplicated multiple times, the same gene can appear in multiple duplicate pairs, often derived from different duplication modes. However, it is often useful to classify genes into unique modes of duplication (e.g., to understand the mechanism behind the expansion of particular gene families). This can be performed with the function *classify_genes()*, which assigns genes to unique duplication modes using the following hierarchy: SD > TD > PD > rTRD > dTRD > DD.

### Calculation of substitution rates per substitution site

Rates of synonymous substitution per substitution site (K_s_), non-synonymous substitutions per substitution site (K_a_), and their ratios (K_a_/K_s_) for each gene pair can be calculated with the function *pairs2kaks()* using all 17 codon-based models of nucleotide substitution from KaKs_Calculator 2.0 (Wang et al. 2010), as implemented in MSA2dist (Ullrich 2023). Peaks in K_s_ distributions, which typically suggest the presence of whole-genome duplication (WGD) events, can be predicted with normal and lognormal mixture modeling using the function *find_ks_peaks()*. However, we advise users to interpret such peaks with caution, as mixture models are prone to overclustering and overfitting (Tiley et al. 2018). Mixture components can further be used to classify gene pairs in age groups with the function *split_pairs_by_peak()*, with age boundaries defined by the intersection point between two different components or by the mean ± *N* standard deviations (default = 2, but can be adjusted).

### Data visualization

Graphical functions are available to create publication-ready plots from the output of *doubletrouble* functions. Absolute and relative frequencies of duplicated genes/gene pairs by mode per species can be visualized with the function *plot_duplicate_freqs()* (Fig. 2). Distributions of substitution rates (K_a_, K_s_, and K_a_/K_s_) per species can be visualized with the function *plot_rates_by_species()* (Fig. 3A). Likewise, K_s_ distributions and peaks can be visualized with the function *plot_ks_distro()* and *plot_ks_peaks()*, respectively (Fig. 3B). All plots are created with the ggplot2 system (Wicham 2016), allowing users to customize plot aesthetics (e.g., color palettes, axis labels, font aspects, themes, etc).

**Fig. 2.**
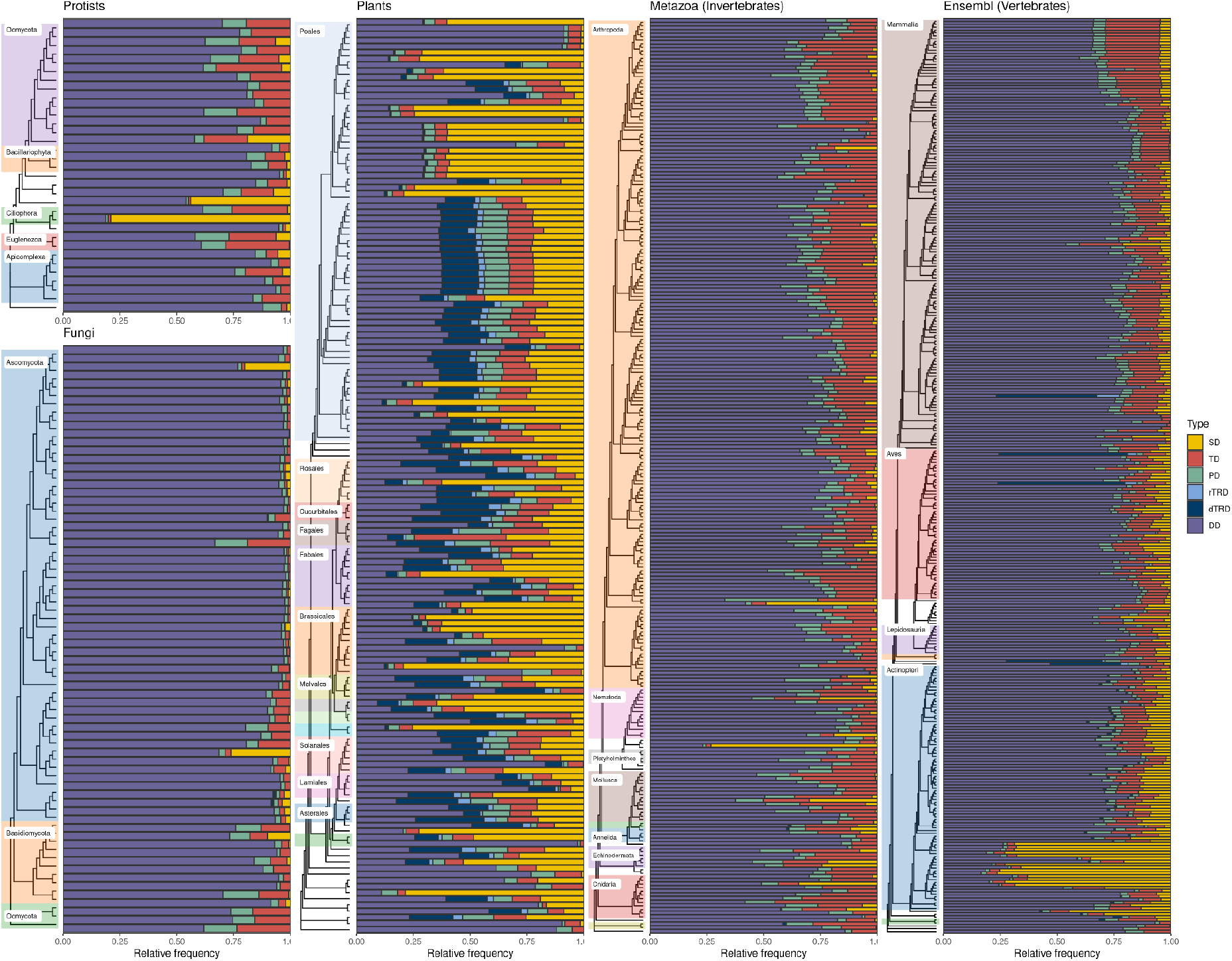
The duplicated gene repertoire in 822 eukaryotic genomes. The figure shows the relative contribution of each duplication mode to the total set of duplicated genes in each genome. Data were obtained from Ensembl and Ensembl instances (Protists, Fungi, Plants, Metazoa, and Vertebrates; see Materials and Methods for details), and duplicates were classified using the ‘full’ scheme (except for Protists, for which an outgroup choice was not trivial). Visual inspection shows that plants are distinctive, with large proportions of segmental duplicates that, in many taxa, represent the major duplication mechanism. Similarly large proportions of segmental duplicates can also be observed in teleost fish (Ensembl Vertebrates) and some other species in other Ensembl instances.

**Fig. 3.**
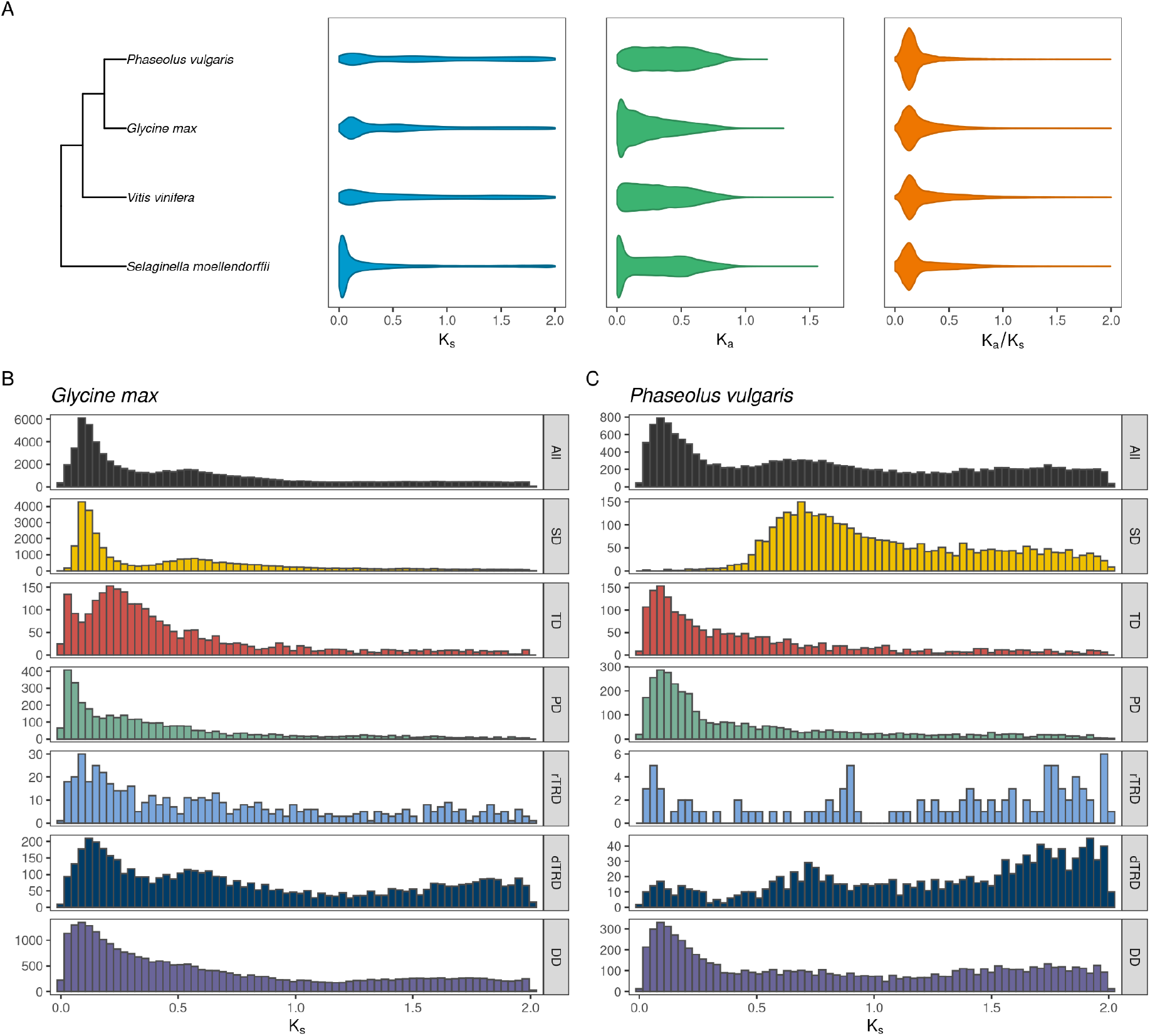
Distribution of substitution rates for selected species. A. Distribution of synonymous substitutions (K_s_), non-synonymous substitutions (K_a_), and K_a_/K_s_ for four plant species in a phylogenetic context, as generated with the function *plot_rates_by_species()*. **B**. K_s_ distribution for all paralogous pairs in the soybean genomes, with facets showing subsets for each duplication mode. and **C**. K_s_ distribution for all paralogous pairs in the common bean genomes, with facets showing subsets for each duplication mode. Figures highlight the importance of visualizing K_s_ distributions for duplicates originating from each duplicated mode separately, especially when looking for evidence of whole-genome duplication events.

### Benchmark data sets

Whole-genome protein sequences and gene annotations were obtained for all species in release 57 of Ensembl Fungi (*N* = 70), Ensembl Protists (*N* = 33), Ensembl Plants (N = 149) and Ensembl Metazoa (N = 253) (Yates et al. 2022), and in release 110 of Ensembl (N = 317) (Martin et al. 2023), leading to a total of 822 genomes across the Eukarya tree of life (Supplementary Table S1). BUSCO scores (Manni et al. 2021) for each species were obtained and visualized using the Bioconductor package cogeqc (Almeida-Silva and Van de Peer 2023). BUSCO genes shared by >90% of the species were aligned with MAFFT (Katoh and Standley 2013), and multiple sequence alignments were concatenated and trimmed to remove alignment columns with >50% gaps. Filtered supermatrices were used for phylogeny inference with IQ-TREE2 (Minh et al. 2020). Oomycetes, red algae, *Giardia lamblia, Mnemiopsis leidyi*, and *Saccharomyces cerevisiae* were used as outgroups for Ensembl Fungi, Ensembl Plants, Ensembl Protists, Ensembl Metazoa, and Ensembl, respectively (Supplementary Text S1).

## Results and Discussion

### Runtime benchmark

To assess doubletrouble’s performance, we classified duplicate pairs with *classify_gene_pairs()* in the genomes of six model organisms, namely budding yeast (*Saccharomyces cerevisiae)*, humans (*Homo sapiens*), fruit fly (*Drosophila melanogaster*), zebrafish (*Danio rerio*), worm (*Caenorhabditis elegans*), and thale cress (*Arabidopsis thaliana*) (Supplementary Text S5). On an Ubuntu 22.04 laptop with an Intel i5-1345U processor (4.7 GHz; 16 GB RAM), duplicate classification took from 0.21 seconds (*S. cerevisiae)* to 5.79 seconds (*D. rerio*), revealing that such task is efficient and can be easily performed on large-scale genomic data sets (Fig. 1B). Next, we calculated substitution rates for all duplicated gene pairs in the *S. cerevisiae* genome (N = 3588). Using a single thread, the function *pairs2kaks()* took 30 minutes to complete (1.98 pairs/second; Fig. 1C), which includes the time to perform a pairwise alignment and calculate substitution rates for each gene pair. When parallelization was enabled with four and eight threads, calculations ran in 7.5 minutes (7.91 pairs/second) and 5 minutes (11.86 pairs/second), respectively (Fig. 1C).

### Use case 1: The duplicated gene repertoire across the Eukarya tree of life

Using genomic data for 822 species in Ensembl and Ensembl Genomes (Fungi, Protists, Plants, and Metazoa), we identified duplicated gene pairs and classified them by duplication mode using the ‘full*’* classification scheme (Supplementary Text S2). For Ensembl Protists, however, we used the ‘standard’ scheme, because protists are not a monophyletic group, hindering the choice of suitable outgroups to detect transposed duplicates. Expectedly, we observed a much larger proportion of SD-derived genes in plants as compared to all other taxa (Fig. 2), which is likely due to pervasive WGD events in the plant tree of life (Van de Peer et al. 2017). In animals, fungi, and protists, tandem duplication is the main source of novel genes. However, most of the duplicates in these taxa are classified as dispersed duplicates, indicating that very little is known about their modes of duplication.

Although large-scale duplications are rare in non-plant organisms, we observed a large proportion of segmental duplicates in some taxa and in independent species (Supplementary Table S2). Importantly, we found weak or no association between the percentages of segmental duplicates and genome completeness in this data set, indicating that the high proportion of segmental duplicates is due to unique evolutionary aspects of some taxa (Supplementary Figure S1; Supplementary Text S4). For instance, most of the duplicated genes in teleost fish originated from segmental duplications (Fig. 2), which is line with established evidence for a fish-specific WGD during the evolution of vertebrates (Amores et al. 1998; Meyer and Van De Peer 2005). Likewise, in the fungal tree of life, large proportions of SD-derived genes are observed in *Saccharomyces cerevisiae* (26.2%) and *Fusarium oxysporum* (20.2%). The former is a paleopolyploid with a genome duplication resulting from an allopolyploidization (Wolfe and Shields 1997; Marcet-Houben and Gabaldón 2015; Escalera-Fanjul et al. 2019), while the latter contains horizontally transferred lineage-specific chromosomes in its genome (Ma et al. 2010; Zhang et al. 2020). Other taxa with high proportions of segmental duplicates include the sea anemone *Actinia equina*, the unicellular eukaryote *Paramecium tetraurelia*, the rotifer *Adineta vaga*, and the barnacle *Amphibalanus amphitrite*, all of which with a suggested history of polyploidization (Aury et al. 2006; Wilding et al. 2020; Simion et al. 2021; Yuan et al. 2023).

### Use case 2: Whole-genome duplications in legumes

Peaks in K_s_ distributions are typically used as evidence of WGD events (Zwaenepoel et al. 2019; Sensalari et al. 2022). However, commonly employed mixture modeling techniques can lead to overclustering (*i*.*e*., prediction of more peaks than the actual number) (Tiley et al. 2018; Zwaenepoel et al. 2019), and including collinearity information can result in more accurate predictions of WGD events (Zwaenepoel et al. 2019; Roelofs et al. 2020). To demonstrate this, we used the function *pairs2kaks()* to calculate substitution rates for all duplicate pairs in the genomes of soybean (*Glycine max*) and common bean (*Phaseolus vulgaris*) (Supplementary Text S3). When visualizing K_s_ distributions for all duplicates, two peaks appear to be present at around 0.1 and 0.6 in both species (Fig. 3B and 3C). However, visualizing the distributions with duplicates classified by mode reveals that, among segmental duplicates, such two peaks can only be observed for soybean, while only the more ancient peak can be observed for common bean (Fig. 3B and 3C). Two WGD events have been dated in the common ancestor of all legumes and the *Glycine* genus, at around 58 and 13 million years ago, respectively (Shoemaker et al. 2006; Fawcett et al. 2009; Koenen et al. 2021). Thus, we expect to detect two peaks and one peak in the K_s_ distributions of soybean and common bean, respectively. Here, we demonstrate that one could mispredict an extra peak for common bean without collinearity information included, highlighting its importance in the prediction of WGD events.

### A web application towards FAIR data principles

To ensure the data generated in this study are FAIR (Findable, Accessible, Interoperable, and Reusable), we developed *doubletroubledb*, a user-friendly web application where users can explore frequencies of duplicated genes for all 822 eukaryote species (see Materials and Methods for details). The web application also works as an interface to download tabular data files with duplicated genes and gene pairs for each species. The application is accessible at https://almeidasilvaf.github.io/doubletroubledb, and the source code is accessible at https://github.com/almeidasilvaf/doubletroubledb.

## Conclusion

*doubletrouble* is an R/Bioconductor package that can be used to identify and classify duplicated genes by duplication mode, calculate substitution rates for gene pairs, and create publication-ready figures. This package should help in exploring the duplication landscape in different species, and the data sets generated in this study will be an important resource for researchers studying the evolution of gene duplication in eukaryotes.

## Acknowledgements

YVdP acknowledges funding from the European Research Council (ERC) under the European Union’s Horizon 2020 research and innovation program (No. 833522). YVdP and FA-S acknowledge funding from Ghent University (Methusalem funding, BOF.MET.2021.0005.01).

## Competing interests

None declared.

## Data availability

To ensure full reproducibility, all code and data used in this manuscript are available in a GitHub repository at https://github.com/almeidasilvaf/doubletrouble_paper, and the code used in benchmarks are available in Supplementary Texts.

## Notes

https://almeidasilvaf.github.io/doubletroubledb

https://github.com/almeidasilvaf/doubletrouble_paper

### Competing Interest Statement

The authors have declared no competing interest.

