## Supplementary Figures for "*doubletrouble:* an R/Bioconductor package for the identification, classification, and analysis of gene and genome duplications"

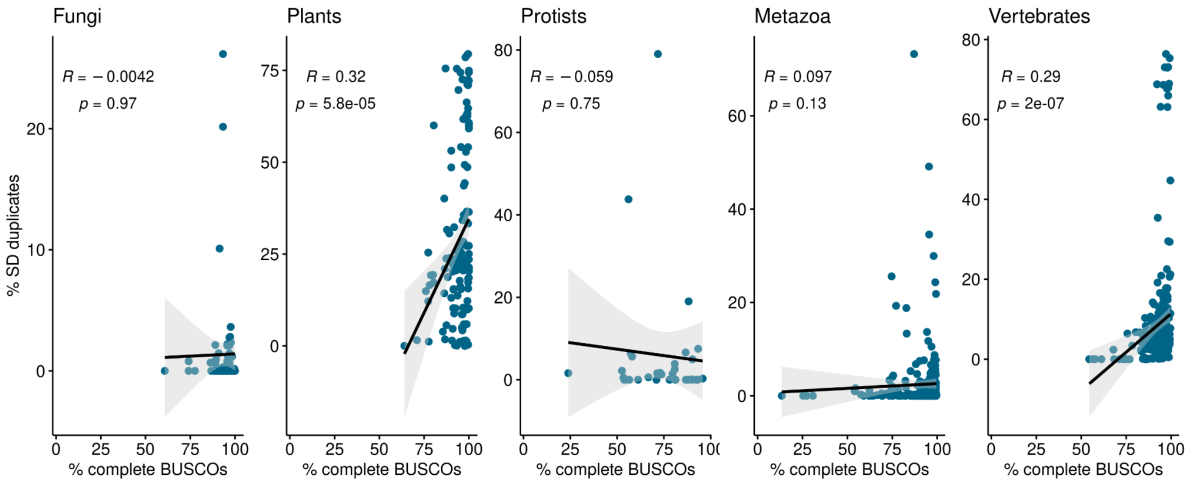


**Fig. 1. Association between the percentage of segmental duplicates and the percentage of complete BUSCOs.** Spearman correlation coefficients and P-values are indicated in each plot. We observed weak or no association between the two variables.
