## Supplementary Texts for "*doubletrouble:* an R/Bioconductor package for the identification, classification, and analysis of gene and genome duplications"

### Supplementary Text S1: Obtaining species trees for Ensembl instances

***Fabricio Almeida-Silva***[1] ***and Yves Van de Peer***[1]

[1]VIB-UGent Center for Plant Systems Biology, Ghent University, Ghent, Belgium

**27 February 2024**

### 1 Introduction

Here, we will describe the code to obtain a species tree for each Ensembl instance using BUSCO genes.

```
library(here)
## here() starts at /home/faalm/Dropbox/package_benchmarks/doubletrouble_paper
library(tidyverse)
## -- Attaching core tidyverse packages ------------------------ tidyverse 2.0.0 --
## v dplyr     1.1.4     v readr     2.1.5
## v forcats   1.0.0     v stringr   1.5.1
## v ggplot2   3.4.4     v tibble    3.2.1
## v lubridate 1.9.3     v tidyr     1.3.1
## v purrr     1.0.2
## -- Conflicts ------------------------------------------ tidyverse_conflicts() --
## x dplyr::filter() masks stats::filter()
## x dplyr::lag()    masks stats::lag()
## i Use the conflicted package (<http://conflicted.r-lib.org/>) to force all conflicts to become errors
library(biomaRt)
library(Herper)
## Loading required package: reticulate
##
## Attaching package: 'Herper'
##
## The following object is masked from 'package:reticulate':
##
##     conda_search
library(taxize)
library(Biostrings)
## Loading required package: BiocGenerics
##
## Attaching package: 'BiocGenerics'
##
## The following objects are masked from 'package:lubridate':
##
##     intersect, setdiff, union
##
## The following objects are masked from 'package:dplyr':
##
##     combine, intersect, setdiff, union
##
## The following objects are masked from 'package:stats':
##
##     IQR, mad, sd, var, xtabs
##
## The following objects are masked from 'package:base':
##
##     anyDuplicated, aperm, append, as.data.frame, basename, cbind,
##     colnames, dirname, do.call, duplicated, eval, evalq, Filter, Find,
##     get, grep, grepl, intersect, is.unsorted, lapply, Map, mapply,
##     match, mget, order, paste, pmax, pmax.int, pmin, pmin.int,
```

```
##     Position, rank, rbind, Reduce, rownames, sapply, setdiff, sort,
##     table, tapply, union, unique, unsplit, which.max, which.min
##
## Loading required package: S4Vectors
## Loading required package: stats4
##
## Attaching package: 'S4Vectors'
##
## The following objects are masked from 'package:lubridate':
##
##     second, second<-
##
## The following objects are masked from 'package:dplyr':
##
##     first, rename
##
## The following object is masked from 'package:tidyr':
##
##     expand
##
## The following object is masked from 'package:utils':
##
##     findMatches
##
## The following objects are masked from 'package:base':
##
##     expand.grid, I, unname
##
## Loading required package: IRanges
##
## Attaching package: 'IRanges'
##
## The following object is masked from 'package:lubridate':
##
##     %within%
##
## The following objects are masked from 'package:dplyr':
##
##     collapse, desc, slice
##
## The following object is masked from 'package:purrr':
##
##     reduce
##
## Loading required package: XVector
##
## Attaching package: 'XVector'
##
## The following object is masked from 'package:purrr':
##
##     compact
```

```
##
## Loading required package: GenomeInfoDb
##
## Attaching package: 'Biostrings'
##
## The following object is masked from 'package:base':
##
##     strsplit
library(cogeqc)

set.seed(123) # for reproducibility
options(timeout = 1e6) # to allow download of big files

source(here("code", "utils.R"))
source(here("code", "utils_busco_phylogeny.R"))
```

### 2 Summary stats

To start with, let's get the number of species for each instance:

```
# Get number of species in Ensembl Genomes
instances <- c("fungi_mart", "plants_mart", "metazoa_mart", "protists_mart")
nspecies_ensemblgenomes <- unlist(lapply(instances, function(x) {
    return(nrow(listDatasets(useEnsemblGenomes(biomart = x))))
}))

# Get number of species in Ensembl
nspecies_ensembl <- nrow(listDatasets(useEnsembl(biomart = "genes")))

# Combine summary stats onto a data frame
nspecies_all <- data.frame(
    instance = c(gsub("_mart", "", instances), "ensembl"),
    n_genes = c(nspecies_ensemblgenomes, nspecies_ensembl)
)

nspecies_all
##   instance n_genes
## 1    fungi      70
## 2   plants     151
## 3  metazoa     280
## 4 protists      33
## 5  ensembl     214
```

### 3 Getting species metadata

Now, let's get species metadata for each Ensembl instance.

```r
# Metadata column names
col_names <- c(
    "name", "species", "division", "taxonomy_id", "assembly",
    "assembly_accession", "genebuild", "variation", "microarray", "pan_compara",
    "peptide_compara", "genome_alignments", "other_alignments", "core_db",
    "species_id"
)

to_remove <- c(
    "variation", "microarray", "pan_compara", "peptide_compara",
    "genome_alignments", "other_alignments", "core_db", "species_id"
)

# Ensembl Fungi
metadata_fungi <- read_tsv(
    "http://ftp.ebi.ac.uk/ensemblgenomes/pub/release-57/fungi/species_EnsemblFungi.txt",
    col_names = col_names, skip = 1, col_select = 1:15, show_col_types = FALSE
) |>
    dplyr::filter(!startsWith(core_db, "fungi_")) |>
    dplyr::select(!any_of(to_remove)) |>
    as.data.frame()

metadata_fungi <- cbind(
    metadata_fungi,
    classification(metadata_fungi$taxonomy_id, db = "ncbi") |>
        format_classification()
)

# Ensembl Plants
metadata_plants <- read_tsv(
    "http://ftp.ebi.ac.uk/ensemblgenomes/pub/release-57/plants/species_EnsemblPlants.txt",
    col_names = col_names, skip = 1, col_select = 1:15, show_col_types = FALSE
) |>
    dplyr::filter(species != "triticum_aestivum_kariega") |>
    dplyr::select(!any_of(to_remove)) |>
    as.data.frame()

metadata_plants <- cbind(
    metadata_plants,
    classification(metadata_plants$taxonomy_id, db = "ncbi") |>
        format_classification()
)

# Ensembl Metazoa
metadata_metazoa <- read_tsv(
    "http://ftp.ebi.ac.uk/ensemblgenomes/pub/release-57/metazoa/species_EnsemblMetazoa.txt",
    col_names = col_names, skip = 1, col_select = 1:15, show_col_types = FALSE
) |>
    dplyr::filter(!startsWith(core_db, "metazoa_")) |>
    dplyr::select(!any_of(to_remove)) |>
    as.data.frame()
```

```
metadata_metazoa <- cbind(
    metadata_metazoa,
    classification(metadata_metazoa$taxonomy_id, db = "ncbi") |>
        format_classification()
)

# Ensembl Protists
metadata_protists <- read_tsv(
    "http://ftp.ebi.ac.uk/ensemblgenomes/pub/release-57/protists/species_EnsemblProtists.txt",
    col_names = col_names, skip = 1, col_select = 1:15, show_col_types = FALSE
) |>
    dplyr::filter(!startsWith(core_db, "protists_")) |>
    dplyr::select(!any_of(to_remove)) |>
    as.data.frame()

metadata_protists <- cbind(
    metadata_protists,
    classification(metadata_protists$taxonomy_id, db = "ncbi") |>
        format_classification()
)

# Ensembl
metadata_ensembl <- read_tsv(
    "https://ftp.ensembl.org/pub/release-110/species_EnsemblVertebrates.txt",
    col_names = col_names, skip = 1, col_select = 1:15, show_col_types = FALSE
) |>
    dplyr::select(!any_of(to_remove)) |>
    as.data.frame()

metadata_ensembl <- cbind(
    metadata_ensembl,
    classification(metadata_ensembl$taxonomy_id, db = "ncbi") |>
        format_classification()
)

# Combining all metadata data frames into a list and saving it
metadata_all <- list(
    fungi = metadata_fungi,
    plants = metadata_plants,
    metazoa = metadata_metazoa,
    protists = metadata_protists,
    ensembl = metadata_ensembl
)

save(
    metadata_all, compress = "xz",
    file = here("products", "result_files", "metadata_all.rda")
)
```

### 4 BUSCO-guided phylogeny inference

Here, for each Ensembl instance, we infer a species tree using the following workflow:

1. Run BUSCO in protein mode with **cogeqc**, using translated sequences for primary transcripts as input;
2. Get the sequences of the identified complete BUSCOs that are shared across all species;
3. Perform a multiple sequence alignment for each BUSCO gene family.
4. Trim the alignments to remove columns with >50% of gaps.
5. Infer a phylogeny with IQ-TREE2.

To start with, we will use the Bioconductor package **Herper** to create a Conda environment containing BUSCO and all its dependencies. Then, we will use this environment to run BUSCO from the R session.

```
# Create Conda environment with BUSCO
my_miniconda <- "~/"

conda <- install_CondaTools(
    tools = "busco==5.5.0",
    env = "busco_env",
    pathToMiniConda = my_miniconda
)
```

#### 4.1 Obtaining BUSCO sequences

To obtain sequences for BUSCO genes, we will run BUSCO in protein mode using the R/Bioconductor package **cogeqc**. Then, we will read the sequences for complete, single-copy BUSCOs, and keep only BUSCO genes that are shared by a certain % of the species. Ideally, this cut-off should be 100% of conservation (i.e., the BUSCO gene is found in all species), but it can be relaxed for some clades.

##### 4.1.1 Ensembl Fungi

Here, we will obtain BUSCO genes for Ensembl Fungi species using the following parameters:

1. Lineage: **eukaryota_odb10**
2. Conservation: 100%

```
# Download whole-genome protein sequences to a directory sequences
busco_fungi <- file.path("~/Downloads/busco_fungi")
seq_fungi <- file.path(busco_fungi, "seqs")
if(!dir.exists(seq_fungi)) { dir.create(seq_fungi, recursive = TRUE) }

download_filtered_proteomes(metadata_all$fungi, "fungi", seq_fungi)

# Run BUSCO in `protein` mode
with_CondaEnv(
    "busco_env",
    cogeqc::run_busco(
        sequence = seq_fungi,
        outlabel = "ensembl_fungi",
```

```
        mode = "protein",
        lineage = "eukaryota_odb10",
        outpath = busco_fungi,
        threads = 3,
        download_path = busco_fungi

    ),
    pathToMiniConda = my_miniconda
)

outdir <- file.path(busco_fungi, "ensembl_fungi")
fungi_busco_seqs <- read_busco_sequences(outdir, verbose = TRUE)
```

Saving BUSCO sequences:

```
# Save list of AAStringSet objects with conserved BUSCO sequences
save(
    fungi_busco_seqs, compress = "xz",
    file = here("products", "result_files", "busco_seqs", "fungi_busco_seqs.rda")
)
```

##### 4.1.2 Ensembl Plants

Here, we will use the lineage data set **eukaryota_odb10**. We could use **viridiplantae_odb10**, but there are 3 Rhodophyta species (*Chondrus crispus*, *Galdieria sulphuraria*, and *Cyanidioschyzon merolae*). Because none of the BUSCO genes were shared by all species, we selected genes shared by >60% of the species, and then manually selected BUSCO genes in a way that all species are included. This was required because some taxa (in particular *Triticum* species) had very few BUSCO genes.

```
# Download whole-genome protein sequences to a directory sequences
busco_plants <- file.path("~/Downloads/busco_plants")
seq_plants <- file.path(busco_plants, "seqs")
if(!dir.exists(seq_plants)) { dir.create(seq_plants, recursive = TRUE) }

download_filtered_proteomes(metadata_all$plants, "plants", seq_plants)

# Run BUSCO in `protein` mode
with_CondaEnv(
    "busco_env",
    cogeqc::run_busco(
        sequence = seq_plants,
        outlabel = "ensemblplants",
        mode = "protein",
        lineage = "eukaryota_odb10",
        outpath = busco_plants,
        threads = 4,
        download_path = busco_plants

    ),
    pathToMiniConda = my_miniconda
```

```
    )

    # Read sequences of BUSCOs preserved in >=60% of the species
    outdir <- file.path(busco_plants, "ensemblplants")
    plants_busco_seqs <- read_busco_sequences(outdir, conservation_freq = 0.6)

    # Select 10 BUSCO genes so that all species are represented
    plants_busco_pav <- get_busco_pav(plants_busco_seqs)

    #' The following code was used to manually select BUSCOs in a way that
    #' all species are represented

    #> ht <- ComplexHeatmap::Heatmap(plants_busco_pav)
    #> ht <- ComplexHeatmap::draw(ht)
    #> InteractiveComplexHeatmap::htShiny(ht)

    # Create a vector of selected BUSCOs
    selected_buscos <- c(
        "549762at2759", "1003258at2759", "1247641at2759",
        "1200489at2759", "1398309at2759", "1346432at2759",
        "1266231at2759", "1094121at2759", "1421503at2759",
        "664730at2759", "1405073at2759", "450058at2759",
        "865202at2759", "901894at2759", "1450538at2759",
        "1284731at2759"
    )

    # Subset sequences to keep only selected BUSCOs
    plants_busco_seqs <- plants_busco_seqs[selected_buscos]
```

Saving BUSCO sequences:

```
# Save list of AAStringSet objects with conserved BUSCO sequences
save(
    plants_busco_seqs, compress = "xz",
    file = here("products", "result_files", "busco_seqs", "plants_busco_seqs.rda")
)
```

##### 4.1.3 Ensembl Protists

Here, we will obtain BUSCO genes for Ensembl Protists species using the following parameters:

1. Lineage: **eukaryota_odb10**
2. Conservation: 100%

```
# Download whole-genome protein sequences to a directory sequences
busco_protists <- file.path("~/Downloads/busco_protists")
seq_protists <- file.path(busco_protists, "seqs")
if(!dir.exists(seq_protists)) { dir.create(seq_protists, recursive = TRUE) }

download_filtered_proteomes(metadata_all$protists, "protists", seq_protists)
```

```r
# Run BUSCO in `protein` mode
with_CondaEnv(
    "busco_env",
    cogeqc::run_busco(
        sequence = seq_protists,
        outlabel = "ensemblprotists",
        mode = "protein",
        lineage = "eukaryota_odb10",
        outpath = busco_protists,
        threads = 4,
        download_path = busco_protists

    ),
    pathToMiniConda = my_miniconda
)

# Read sequences of BUSCOs preserved in >=60% of the species
outdir <- file.path(busco_protists, "ensemblprotists")
protists_busco_seqs <- read_busco_sequences(outdir, verbose = TRUE)
```

Saving BUSCO sequences:

```r
# Save list of AAStringSet objects with conserved BUSCO sequences
save(
    protists_busco_seqs, compress = "xz",
    file = here("products", "result_files", "busco_seqs", "protists_busco_seqs.rda")
)
```

##### 4.1.4 Ensembl Metazoa

For the Metazoa instance, we used the *metazoa_odb10* lineage data set.

```r
# Download whole-genome protein sequences to a directory sequences
busco_metazoa <- file.path("~/Downloads/busco_metazoa")
seq_metazoa <- file.path(busco_metazoa, "seqs")
if(!dir.exists(seq_metazoa)) { dir.create(seq_metazoa, recursive = TRUE) }

download_filtered_proteomes(metadata_all$metazoa, "metazoa", seq_metazoa)

# Run BUSCO in `protein` mode
with_CondaEnv(
    "busco_env",
    cogeqc::run_busco(
        sequence = seq_metazoa,
        outlabel = "ensemblmetazoa",
        mode = "protein",
        lineage = "metazoa_odb10",
        outpath = busco_metazoa,
        threads = 4,
        download_path = busco_metazoa
```

```
        ),
        pathToMiniConda = my_miniconda
    )

    # Read sequences of BUSCOs preserved in >=60% of the species
    outdir <- file.path(busco_metazoa, "ensemblmetazoa")
    metazoa_busco_seqs <- read_busco_sequences(outdir, conservation_freq = 0.9)

    # Select 10 BUSCO genes so that all species are represented
    metazoa_busco_pav <- get_busco_pav(metazoa_busco_seqs)

    #' The following code was used to manually select BUSCOs in a way that
    #' all species are represented

    #> ht <- ComplexHeatmap::Heatmap(metazoa_busco_pav)
    #> ht <- ComplexHeatmap::draw(ht)
    #> InteractiveComplexHeatmap::htShiny(ht)

    # Create a vector of selected BUSCOs
    selected_buscos <- c(
        "351226at33208", "135294at33208",
        "517525at33208", "501396at33208",
        "464987at33208", "443518at33208",
        "495100at33208", "335107at33208",
        "454911at33208", "134492at33208"
    )

    # Subset sequences to keep only selected BUSCOs
    metazoa_busco_seqs <- metazoa_busco_seqs[selected_buscos]
```

Saving BUSCO sequences:

```
    # Save list of AAStringSet objects with conserved BUSCO sequences
    save(
        metazoa_busco_seqs, compress = "xz",
        file = here("products", "result_files", "busco_seqs", "metazoa_busco_seqs.rda")
    )
```

##### 4.1.5 Ensembl Vertebrates

Here, because there are 3 non-vertebrate species (*C. elegans*, *D. melanogaster*, and *S. cerevisiae*), we will use the lineage data set **eukaryota_odb10**.

```
    # Download whole-genome protein sequences to a directory sequences
    busco_vertebrates <- file.path("~/Downloads/busco_vertebrates")
    seq_vertebrates <- file.path(busco_vertebrates, "seqs")
    if(!dir.exists(seq_vertebrates)) { dir.create(seq_vertebrates, recursive = TRUE) }

    download_filtered_proteomes(metadata_all$ensembl, "ensembl", seq_vertebrates)
```

```r
# Run BUSCO in `protein` mode
with_CondaEnv(
    "busco_env",
    cogeqc::run_busco(
        sequence = seq_vertebrates,
        outlabel = "ensemblvertebrates",
        mode = "protein",
        lineage = "eukaryota_odb10",
        outpath = busco_vertebrates,
        threads = 4,
        download_path = busco_vertebrates

    ),
    pathToMiniConda = my_miniconda
)

# Read sequences of BUSCOs preserved in >=90% of the species
outdir <- file.path(busco_vertebrates, "ensemblvertebrates")
vertebrates_busco_seqs <- read_busco_sequences(outdir, conservation_freq = 0.9)

# Select 10 BUSCO genes so that all species are represented
vertebrates_busco_pav <- get_busco_pav(vertebrates_busco_seqs)

#' The following code was used to manually select BUSCOs in a way that
#' all species are represented

#> ht <- ComplexHeatmap::Heatmap(vertebrates_busco_pav)
#> ht <- ComplexHeatmap::draw(ht)
#> InteractiveComplexHeatmap::htShiny(ht)

# Create a vector of selected BUSCOs
selected_buscos <- c(
    "834694at2759", "551907at2759",
    "491869at2759", "1085752at2759",
    "801857at2759", "1398309at2759",
    "176625at2759", "1324510at2759",
    "1377237at2759", "1085752at2759"
)

# Subset sequences to keep only selected BUSCOs
vertebrates_busco_seqs <- vertebrates_busco_seqs[selected_buscos]
```

Saving BUSCO sequences:

```r
# Save list of AAStringSet objects with conserved BUSCO sequences
save(
    vertebrates_busco_seqs, compress = "xz",
    file = here("products", "result_files", "busco_seqs", "vertebrates_busco_seqs")
)
```

#### 4.2 Tree inference from BUSCO genes

Now, we will infer species trees from MSAs for each family, and from a single concatenated MSA (when possible).

##### 4.2.1 Ensembl Fungi

Performing MSA with MAFFT and trimming the alignment:

```
# Perform MSA with MAFFT
aln_fungi <- align_sequences(busco_seqs_fungi, threads = 4)

# Trim alignment to remove columns with >50% of gaps
aln_fungi_trimmed <- lapply(aln_fungi, trim_alignment, max_gap = 0.5)
```

Now, let's infer a species tree using IQ-TREE2.

```
outgroup <- "aphanomyces.astaci,aphanomyces.invadans,globisporangium.ultimum"
trees_fungi <- infer_species_tree(aln_fungi_trimmed, outgroup, threads = 4)
```

Finally, for comparative reasons, we will also infer a single tree from a concatenated multiple sequence alignment.

```
# Concatenate alignments
aln_fungi_conc <- Reduce(xscat, aln_fungi_trimmed)
names(aln_fungi_conc) <- names(aln_fungi_trimmed[[1]])

# Infer tree from concatenated alignment
tree_fungi_conc <- infer_species_tree(
    list(conc = aln_fungi_conc),
    outgroup, threads = 4
)
```

Combining the trees and saving them:

```
# Combine trees
fungi_busco_trees <- c(
    tree_fungi_conc, trees_fungi
)

save(
    fungi_busco_trees, compress = "xz",
    file = here("products", "result_files", "trees", "fungi_busco_trees.rda")
)
```

##### 4.2.2 Ensembl Plants

Here, because no BUSCO gene is present in all species, we will only infer a single tree from concatenated alignments.

```
# Perform MSA with MAFFT
aln_plants <- align_sequences(plants_busco_seqs, threads = 4)
```

```
# Trim alignment to remove columns with >50% of gaps
aln_plants_trimmed <- lapply(aln_plants, trim_alignment, max_gap = 0.5)
```

Finally, let's infer a species tree from a concatenated alignment. As outgroups, we're going to use *Chondrus crispus*, *Galdieria sulphuraria*, and *Cyanidioschyzon merolae*.

```
outgroup <- "chondrus.crispus,galdieria.sulphuraria,cyanidioschyzon.merolae"

# Concatenate alignments
aln_plants_conc <- concatenate_alignments(aln_plants_trimmed)

# Infer tree from concatenated alignment
plants_busco_trees <- infer_species_tree(
    list(conc = aln_plants_conc),
    outgroup, threads = 4
)

# Save tree
save(
    plants_busco_trees, compress = "xz",
    file = here("products", "result_files", "trees", "plants_busco_trees.rda")
)
```

##### 4.2.3 Ensembl Protists

For this instance, two BUSCO genes were conserved across all species, so we will infer trees for each family + a tree from a concatenated alignment.

```
# Perform MSA with MAFFT
aln_protists <- align_sequences(protists_busco_seqs, threads = 4)

# Trim alignment to remove columns with >50% of gaps
aln_protists_trimmed <- lapply(aln_protists, trim_alignment, max_gap = 0.5)
```

Now, let's infer species trees. As outgroup, we will use Fornicata (*Giardia lamblia*) based on this paper.

```
outgroup <- "giardia.lamblia"

# Path 1: a tree per BUSCO gene
protists_trees1 <- infer_species_tree(
    aln_protists_trimmed, outgroup, threads = 4
)

# Path 2: a single tree from a concatenated alignment
protists_trees2 <- infer_species_tree(
    list(conc = concatenate_alignments(aln_protists_trimmed)),
    outgroup, threads = 6
)
```

```
# Combine trees and save them
protists_busco_trees <- c(protists_trees1, protists_trees2)

save(
    protists_busco_trees, compress = "xz",
    file = here("products", "result_files", "trees", "protists_busco_trees.rda")
)
```

However, even though we specified *Giardia lamblia*, IQ-TREE2 placed it as an ingroup. This suggests that, based on our data (BUSCO sequences), *Giardia lamblia* may not be a good outgroup.

Since protists are not actually a real phylogenetic group (not monophyletic), instead of digging deeper into the real phylogeny of the group and searching for a proper outgroup, we will simply use this phylogeny but acknowledging that it may not be completely accurate.

##### 4.2.4 Ensembl Metazoa

For this instance, two BUSCO genes were conserved across all species, so we will infer trees for each family + a tree from a concatenated alignment.

```
# Perform MSA with MAFFT
aln_metazoa <- align_sequences(metazoa_busco_seqs, threads = 4)

# Trim alignment to remove columns with >50% of gaps
aln_metazoa_trimmed <- lapply(aln_metazoa, trim_alignment, max_gap = 0.5)
```

Now, let's infer a species tree. As outgroup, we will use the ctenophore *Mnemiopsis leidyi*.

```
outgroup <- "mnemiopsis.leidyi"

# Get a single tree from a concatenated alignment
metazoa_busco_trees <- infer_species_tree(
    list(conc = concatenate_alignments(aln_metazoa_trimmed)),
    outgroup, threads = 6
)

# Save tree
save(
    metazoa_busco_trees, compress = "xz",
    file = here("products", "result_files", "trees", "metazoa_busco_trees.rda")
)
```

##### 4.2.5 Ensembl Vertebrates

For this instance, no BUSCO gene was conserved in all species. Thus, we will infer a single tree from a concatenated alignment of ten representative BUSCOs.

```
# Perform MSA with MAFFT
aln_vertebrates <- align_sequences(vertebrates_busco_seqs, threads = 4)
```

```
# Trim alignment to remove columns with >50% of gaps
aln_vertebrates_trimmed <- lapply(aln_vertebrates, trim_alignment, max_gap = 0.5)
```

Now, let's infer a species tree. As outgroup, we will use the yeast *Saccharomyces cerevisiae*.

```
outgroup <- "saccharomyces.cerevisiae"

# Get a single tree from a concatenated alignment
vertebrates_busco_trees <- infer_species_tree(
    list(conc = concatenate_alignments(aln_vertebrates_trimmed)),
    outgroup, threads = 6
)

# Save tree
save(
    vertebrates_busco_trees, compress = "xz",
    file = here("products", "result_files", "trees", "vertebrates_busco_trees.rda")
)
```

### 5 Obtaining BUSCO scores

Finally, since we ran BUSCO to obtain single-copy gene families, we will also use BUSCO's output to explore gene space completeness across species in Ensembl instances.

```
# Read BUSCO completeness stats
## Ensembl Fungi
fungi_busco_scores <- read_busco(
    "~/Downloads/busco_fungi/ensembl_fungi"
)

## Ensembl Plants
plants_busco_scores <- read_busco(
    "~/Downloads/busco_plants/ensemblplants"
)

## Ensembl Protists
protists_busco_scores <- read_busco(
    "~/Downloads/busco_protists/ensemblprotists"
)

## Ensembl Metazoa
metazoa_busco_scores <- read_busco(
    "~/Downloads/busco_metazoa/ensemblmetazoa"
)

## Ensembl Vertebrates
vertebrates_busco_scores <- read_busco(
    "~/Downloads/busco_vertebrates/ensemblvertebrates"
)
```

```r
# Save files
save(
    fungi_busco_scores, compress = "xz",
    file = here(
        "products", "result_files", "busco_scores", "fungi_busco_scores.rda"
    )
)

save(
    plants_busco_scores, compress = "xz",
    file = here(
        "products", "result_files", "busco_scores", "plants_busco_scores.rda"
    )
)

save(
    protists_busco_scores, compress = "xz",
    file = here(
        "products", "result_files", "busco_scores", "protists_busco_scores.rda"
    )
)

save(
    metazoa_busco_scores, compress = "xz",
    file = here(
        "products", "result_files", "busco_scores", "metazoa_busco_scores.rda"
    )
)

save(
    vertebrates_busco_scores, compress = "xz",
    file = here(
        "products", "result_files", "busco_scores", "vertebrates_busco_scores.rda"
    )
)
```

### Session info

This document was created under the following conditions:

```
## - Session info ---------------------------------------------------------------
##  setting  value
##  version  R version 4.3.2 (2023-10-31)
##  os       Ubuntu 22.04.3 LTS
##  system   x86_64, linux-gnu
##  ui       X11
##  language (EN)
##  collate  en_US.UTF-8
##  ctype    en_US.UTF-8
```

```
##  tz       Europe/Brussels
##  date     2024-02-27
##  pandoc   3.1.1 @ /usr/lib/rstudio/resources/app/bin/quarto/bin/tools/ (via rmarkdown)
##
## - Packages -------------------------------------------------------------------
##  package          * version     date (UTC) lib source
##  AnnotationDbi      1.64.1      2023-11-03 [1] Bioconductor
##  ape                5.7-1       2023-03-13 [1] CRAN (R 4.3.2)
##  aplot              0.2.2       2023-10-06 [1] CRAN (R 4.3.2)
##  beeswarm           0.4.0       2021-06-01 [1] CRAN (R 4.3.2)
##  Biobase            2.62.0      2023-10-24 [1] Bioconductor
##  BiocFileCache      2.10.1      2023-10-26 [1] Bioconductor
##  BiocGenerics     * 0.48.1      2023-11-01 [1] Bioconductor
##  BiocManager        1.30.22     2023-08-08 [1] CRAN (R 4.3.2)
##  BiocStyle        * 2.30.0      2023-10-24 [1] Bioconductor
##  biomaRt          * 2.58.2      2024-01-30 [1] Bioconductor 3.18 (R 4.3.2)
##  Biostrings       * 2.70.2      2024-01-28 [1] Bioconductor 3.18 (R 4.3.2)
##  bit                4.0.5       2022-11-15 [1] CRAN (R 4.3.2)
##  bit64              4.0.5       2020-08-30 [1] CRAN (R 4.3.2)
##  bitops             1.0-7       2021-04-24 [1] CRAN (R 4.3.2)
##  blob               1.2.4       2023-03-17 [1] CRAN (R 4.3.2)
##  bold               1.3.0       2023-05-02 [1] CRAN (R 4.3.2)
##  bookdown           0.37        2023-12-01 [1] CRAN (R 4.3.2)
##  cachem             1.0.8       2023-05-01 [1] CRAN (R 4.3.2)
##  cli                3.6.2       2023-12-11 [1] CRAN (R 4.3.2)
##  codetools          0.2-19      2023-02-01 [4] CRAN (R 4.2.2)
##  cogeqc           * 1.6.2       2024-01-14 [1] Bioconductor 3.18 (R 4.3.2)
##  colorspace         2.1-0       2023-01-23 [1] CRAN (R 4.3.2)
##  conditionz         0.1.0       2019-04-24 [1] CRAN (R 4.3.2)
##  crayon             1.5.2       2022-09-29 [1] CRAN (R 4.3.2)
##  crul               1.4.0       2023-05-17 [1] CRAN (R 4.3.2)
##  curl               5.2.0       2023-12-08 [1] CRAN (R 4.3.2)
##  data.table         1.15.0      2024-01-30 [1] CRAN (R 4.3.2)
##  DBI                1.2.1       2024-01-12 [1] CRAN (R 4.3.2)
##  dbplyr             2.4.0       2023-10-26 [1] CRAN (R 4.3.2)
##  digest             0.6.34      2024-01-11 [1] CRAN (R 4.3.2)
##  dplyr            * 1.1.4       2023-11-17 [1] CRAN (R 4.3.2)
##  evaluate           0.23        2023-11-01 [1] CRAN (R 4.3.2)
##  fansi              1.0.6       2023-12-08 [1] CRAN (R 4.3.2)
##  fastmap            1.1.1       2023-02-24 [1] CRAN (R 4.3.2)
##  filelock           1.0.3       2023-12-11 [1] CRAN (R 4.3.2)
##  forcats          * 1.0.0       2023-01-29 [1] CRAN (R 4.3.2)
##  foreach            1.5.2       2022-02-02 [1] CRAN (R 4.3.2)
##  fs                 1.6.3       2023-07-20 [1] CRAN (R 4.3.2)
##  generics           0.1.3       2022-07-05 [1] CRAN (R 4.3.2)
##  GenomeInfoDb     * 1.38.6      2024-02-08 [1] Bioconductor 3.18 (R 4.3.2)
##  GenomeInfoDbData   1.2.11      2023-12-21 [1] Bioconductor
##  ggbeeswarm         0.7.2       2023-04-29 [1] CRAN (R 4.3.2)
##  ggfun              0.1.4       2024-01-19 [1] CRAN (R 4.3.2)
##  ggplot2          * 3.4.4       2023-10-12 [1] CRAN (R 4.3.2)
##  ggplotify          0.1.2       2023-08-09 [1] CRAN (R 4.3.2)
```

```
##  ggtree              3.10.0     2023-10-24 [1] Bioconductor
##  glue                1.7.0      2024-01-09 [1] CRAN (R 4.3.2)
##  gridGraphics        0.5-1      2020-12-13 [1] CRAN (R 4.3.2)
##  gtable              0.3.4      2023-08-21 [1] CRAN (R 4.3.2)
##  here              * 1.0.1      2020-12-13 [1] CRAN (R 4.3.2)
##  Herper            * 1.12.0     2023-10-24 [1] Bioconductor
##  hms                 1.1.3      2023-03-21 [1] CRAN (R 4.3.2)
##  htmltools           0.5.7      2023-11-03 [1] CRAN (R 4.3.2)
##  httpcode            0.3.0      2020-04-10 [1] CRAN (R 4.3.2)
##  httr                1.4.7      2023-08-15 [1] CRAN (R 4.3.2)
##  igraph              2.0.1.1    2024-01-30 [1] CRAN (R 4.3.2)
##  IRanges           * 2.36.0     2023-10-24 [1] Bioconductor
##  iterators           1.0.14     2022-02-05 [1] CRAN (R 4.3.2)
##  jsonlite            1.8.8      2023-12-04 [1] CRAN (R 4.3.2)
##  KEGGREST            1.42.0     2023-10-24 [1] Bioconductor
##  knitr               1.45       2023-10-30 [1] CRAN (R 4.3.2)
##  lattice             0.22-5     2023-10-24 [4] CRAN (R 4.3.1)
##  lazyeval            0.2.2      2019-03-15 [1] CRAN (R 4.3.2)
##  lifecycle           1.0.4      2023-11-07 [1] CRAN (R 4.3.2)
##  lubridate         * 1.9.3      2023-09-27 [1] CRAN (R 4.3.2)
##  magrittr            2.0.3      2022-03-30 [1] CRAN (R 4.3.2)
##  Matrix              1.6-3      2023-11-14 [4] CRAN (R 4.3.2)
##  memoise             2.0.1      2021-11-26 [1] CRAN (R 4.3.2)
##  munsell             0.5.0      2018-06-12 [1] CRAN (R 4.3.2)
##  nlme                3.1-163    2023-08-09 [4] CRAN (R 4.3.1)
##  patchwork           1.2.0      2024-01-08 [1] CRAN (R 4.3.2)
##  pillar              1.9.0      2023-03-22 [1] CRAN (R 4.3.2)
##  pkgconfig           2.0.3      2019-09-22 [1] CRAN (R 4.3.2)
##  plyr                1.8.9      2023-10-02 [1] CRAN (R 4.3.2)
##  png                 0.1-8      2022-11-29 [1] CRAN (R 4.3.2)
##  prettyunits         1.2.0      2023-09-24 [1] CRAN (R 4.3.2)
##  progress            1.2.3      2023-12-06 [1] CRAN (R 4.3.2)
##  purrr             * 1.0.2      2023-08-10 [1] CRAN (R 4.3.2)
##  R6                  2.5.1      2021-08-19 [1] CRAN (R 4.3.2)
##  rappdirs            0.3.3      2021-01-31 [1] CRAN (R 4.3.2)
##  Rcpp                1.0.12     2024-01-09 [1] CRAN (R 4.3.2)
##  RCurl               1.98-1.14  2024-01-09 [1] CRAN (R 4.3.2)
##  readr             * 2.1.5      2024-01-10 [1] CRAN (R 4.3.2)
##  reshape2            1.4.4      2020-04-09 [1] CRAN (R 4.3.2)
##  reticulate        * 1.35.0     2024-01-31 [1] CRAN (R 4.3.2)
##  rjson               0.2.21     2022-01-09 [1] CRAN (R 4.3.2)
##  rlang               1.1.3      2024-01-10 [1] CRAN (R 4.3.2)
##  rmarkdown           2.25       2023-09-18 [1] CRAN (R 4.3.2)
##  rprojroot           2.0.4      2023-11-05 [1] CRAN (R 4.3.2)
##  RSQLite             2.3.5      2024-01-21 [1] CRAN (R 4.3.2)
##  rstudioapi          0.15.0     2023-07-07 [1] CRAN (R 4.3.2)
##  S4Vectors         * 0.40.2     2023-11-23 [1] Bioconductor 3.18 (R 4.3.2)
##  scales              1.3.0      2023-11-28 [1] CRAN (R 4.3.2)
##  sessioninfo         1.2.2      2021-12-06 [1] CRAN (R 4.3.2)
##  stringi             1.8.3      2023-12-11 [1] CRAN (R 4.3.2)
##  stringr           * 1.5.1      2023-11-14 [1] CRAN (R 4.3.2)
```

```
##  taxize          * 0.9.100     2022-04-22 [1] CRAN (R 4.3.2)
##  tibble          * 3.2.1       2023-03-20 [1] CRAN (R 4.3.2)
##  tidyr           * 1.3.1       2024-01-24 [1] CRAN (R 4.3.2)
##  tidyselect        1.2.0       2022-10-10 [1] CRAN (R 4.3.2)
##  tidytree          0.4.6       2023-12-12 [1] CRAN (R 4.3.2)
##  tidyverse       * 2.0.0       2023-02-22 [1] CRAN (R 4.3.2)
##  timechange        0.3.0       2024-01-18 [1] CRAN (R 4.3.2)
##  treeio            1.26.0      2023-10-24 [1] Bioconductor
##  tzdb              0.4.0       2023-05-12 [1] CRAN (R 4.3.2)
##  utf8              1.2.4       2023-10-22 [1] CRAN (R 4.3.2)
##  uuid              1.2-0       2024-01-14 [1] CRAN (R 4.3.2)
##  vctrs             0.6.5       2023-12-01 [1] CRAN (R 4.3.2)
##  vipor             0.4.7       2023-12-18 [1] CRAN (R 4.3.2)
##  withr             3.0.0       2024-01-16 [1] CRAN (R 4.3.2)
##  xfun              0.42        2024-02-08 [1] CRAN (R 4.3.2)
##  XML               3.99-0.16.1 2024-01-22 [1] CRAN (R 4.3.2)
##  xml2              1.3.6       2023-12-04 [1] CRAN (R 4.3.2)
##  XVector         * 0.42.0      2023-10-24 [1] Bioconductor
##  yaml              2.3.8       2023-12-11 [1] CRAN (R 4.3.2)
##  yulab.utils       0.1.4       2024-01-28 [1] CRAN (R 4.3.2)
##  zlibbioc          1.48.0      2023-10-24 [1] Bioconductor
##  zoo               1.8-12      2023-04-13 [1] CRAN (R 4.3.2)
##
##  [1] /home/faalm/R/x86_64-pc-linux-gnu-library/4.3
##  [2] /usr/local/lib/R/site-library
##  [3] /usr/lib/R/site-library
##  [4] /usr/lib/R/library
##
## ------------------------------------------------------------------------------
```

### Supplementary Text S2: Identification and classification of duplicated genes in Ensembl and Ensembl Genomes

***Fabricio Almeida-Silva*** [1] ***and Yves Van de Peer*** [1]

[1]VIB-UGent Center for Plant Systems Biology, Ghent University, Ghent, Belgium

**27 February 2024**

**Supplementary Text S2: Identification and classification of duplicated genes in Ensembl and Ensembl Genomes**

### 1 Introduction

Here, we will describe the code to identify and classify duplicated genes in Ensembl and Ensembl Genomes species using the Bioconductor package *doubletrouble*.

```
library(syntenet)
library(doubletrouble)
library(biomaRt)
library(here)
## here() starts at /home/faalm/Dropbox/package_benchmarks/doubletrouble_paper
library(tidyverse)
## -- Attaching core tidyverse packages ------------------------ tidyverse 2.0.0 --
## v dplyr     1.1.4     v readr     2.1.5
## v forcats   1.0.0     v stringr   1.5.1
## v ggplot2   3.4.4     v tibble    3.2.1
## v lubridate 1.9.3     v tidyr     1.3.1
## v purrr     1.0.2
## -- Conflicts ------------------------------------------ tidyverse_conflicts() --
## x dplyr::filter() masks stats::filter()
## x dplyr::lag()    masks stats::lag()
## x dplyr::select() masks biomaRt::select()
## i Use the conflicted package (<http://conflicted.r-lib.org/>) to force all conflicts to become errors

set.seed(123) # for reproducibility
options(timeout = 1e10) # to allow download of big files

# Load helper functions
source(here("code", "utils.R"))
```

### 2 Data loading: species trees and metadata

Here, we will load the data frames of species metadata and `phylo` objects with species trees for each Ensembl instance.

```
# Load metadata
load(here("products", "result_files", "metadata_all.rda"))
names(metadata_all)
## [1] "fungi"    "plants"   "metazoa"  "protists" "ensembl"

# Load trees
load(here("products", "result_files", "trees", "fungi_busco_trees.rda"))
load(here("products", "result_files", "trees", "plants_busco_trees.rda"))
load(here("products", "result_files", "trees", "metazoa_busco_trees.rda"))
load(here("products", "result_files", "trees", "vertebrates_busco_trees.rda"))
load(here("products", "result_files", "trees", "protists_busco_trees.rda"))
```

### 3 Identification and classification of duplicated genes in Ensembl and Ensembl Genomes

Now, let's use *doubletrouble* to identify duplicated genes and classify them using the Ensembl and Ensembl Genomes data sets. Here, to avoid code repetition and optimize memory usage, we will use the wrapper function `ensembl2duplicates()` (in the file *utils.R*). For each species in the metadata data frame, this function:

1. Retrieves whole-genome protein sequences (`AAStringSet`) and gene annotation (`GRanges`) from an Ensembl instance;
2. Filters the `AAStringSet` object to include only the longest protein for each gene (i.e., the translated sequence of the primary transcript);
3. Processes the sequences and annotation with `syntenet::process_input()`;
4. Identifies the paranome with `syntenet::run_diamond()` + identifies orthologs between the query species and an outgroup (optional);
5. Classifies paralogs by duplication modes.

#### 3.1 Ensembl Fungi

First, let's create a data frame with species and their outgroups. Here, we will use the basidiomycete *Cryptococcus neoformans* as outgroup for Ascomycota species, and the oomycete *Aphanomyces astaci* as outgroup for Basidiomycota species.

```
col_dir <- here("products", "result_files", "collinearity", "fungi")
if(!dir.exists(col_dir)) { dir.create(col_dir, recursive = TRUE) }

# Create data frame of query species and outgroup
fungi_outgroups <- metadata_all$fungi |>
    filter(phylum != "Oomycota") |>
    mutate(
        query = species,
        outgroup = case_when(
            phylum == "Ascomycota" ~ "cryptococcus_neoformans",
            TRUE ~ "aphanomyces_astaci"
        )
    ) |>
    select(query, outgroup)

# Identifying and classifying paralogs
fungi_duplicates <- ensembl2duplicates(
    metadata_all$fungi, ensembl = "fungi",
    outgroups = fungi_outgroups,
    collinearity_dir = col_dir
)

# Classify genes into unique duplication modes
fungi_duplicates_unique <- classify_genes(fungi_duplicates)

# Save classification results
```

```
## Duplicate pairs
save(
    fungi_duplicates,
    file = here("products", "result_files", "fungi_duplicates.rda"),
    compress = "xz"
)

## Duplicated genes (unique duplication modes)
save(
    fungi_duplicates_unique,
    file = here("products", "result_files", "fungi_duplicates_unique.rda"),
    compress = "xz"
)
```

#### 3.2 Ensembl Protists

Since protists are not a real (i.e., monophyletic) group, defining an outgroup species is very problematic. For this reason, we will classify duplicates using the *standard* classification scheme here.

```
col_dir <- here("products", "result_files", "collinearity", "protists")
if(!dir.exists(col_dir)) { dir.create(col_dir, recursive = TRUE) }

# Identifying and classifying paralogs
protists_duplicates <- ensembl2duplicates(
    metadata_all$protists, ensembl = "protists", collinearity_dir = col_dir
)

# Classify genes into unique duplication modes
protists_duplicates_unique <- classify_genes(protists_duplicates)

# Save classification results
## Duplicate pairs
save(
    protists_duplicates,
    file = here("products", "result_files", "protists_duplicates.rda"),
    compress = "xz"
)

## Duplicated genes (unique duplication modes)
save(
    protists_duplicates_unique,
    file = here("products", "result_files", "protists_duplicates_unique.rda"),
    compress = "xz"
)
```

#### 3.3 Ensembl Plants

Here, we will use different outgroups for different branches of the tree. The clades and outgroups are:

1. Angiosperms: *Amborella trichopoda* as outgroup.
2. *Amborella trichopoda* and *Nymphaea colorata*: *Chara braunii* as outgroup.
3. *Selaginella moellendorffii*, *Chara braunii*, *Marchantia polymorpha*, *Physcomitrium patens*: *Chlamydomonas reinhardtii* as outgroup.
4. *Chlamydomonas reinhardtii* and *Ostreococcus lucimarinus*: *Galdieria sulphuraria* as outgroup
5. Rhodophyta algae: no outgroup.

```r
# Create data frame of query species and outgroup
angiosperms <- metadata_all$plants |>
    filter(
        phylum == "Streptophyta",
        !order %in% c(
            "Charales", "Selaginellales", "Funariales",
            "Marchantiales", "Nymphaeales"
        )
    ) |>
    pull(species)

ana <- c("amborella_trichopoda", "nymphaea_colorata")
bryophytes <- c(
    "selaginella_moellendorffii", "chara_braunii",
    "marchantia_polymorpha", "physcomitrium_patens"
)
chlorophyta <- c("chlamydomonas_reinhardtii", "ostreococcus_lucimarinus")

plants_outgroups <- metadata_all$plants |>
    filter(phylum != "Rhodophyta") |>
    mutate(
        query = species,
        outgroup = case_when(
            species %in% angiosperms ~ "amborella_trichopoda",
            species %in% ana ~ "chara_braunii",
            species %in% bryophytes ~ "chlamydomonas_reinhardtii",
            species %in% chlorophyta ~ "galdieria_sulphuraria"
        )
    ) |>
    select(query, outgroup)
```

Identifying and classifying duplicates:

```r
col_dir <- here("products", "result_files", "collinearity", "plants")
if(!dir.exists(col_dir)) { dir.create(col_dir, recursive = TRUE) }

# Identifying and classifying paralogs
plants_duplicates <- ensembl2duplicates(
    metadata_all$plants, ensembl = "plants",
    outgroups = plants_outgroups,
    collinearity_dir = col_dir,
    threads = 4
```

```
)

# Classify genes into unique duplication modes
plants_duplicates_unique <- classify_genes(plants_duplicates)

# Save classification results
## Duplicate pairs
save(
    plants_duplicates,
    file = here("products", "result_files", "plants_duplicates.rda"),
    compress = "xz"
)

## Duplicated genes (unique duplication modes)
save(
    plants_duplicates_unique,
    file = here("products", "result_files", "plants_duplicates_unique.rda"),
    compress = "xz"
)
```

#### 3.4 Ensembl Metazoa

Here, we will use different outgroups for different branches of the tree. The clades and outgroups are:

1. Arthropoda: *Hypsibius exemplaris* (Tardigrada) as outgroup.
2. Tardigrada, Acanthocephala, and Rotifera: *Brugia malayi* (Nematoda) as outgroup.
3. Nematoda: *Priapulus caudatus* (Priapulida) as outgroup
4. Priapulida, Echinodermata, Chordata, and Hemichordata: *Hofstenia miamia* (Xenacoelomorpha) as outgroup.
5. Xenacoelomorpha: *Actinia tenebrosa* (Cnidaria) as outgroup.
6. Cnidaria and Placozoa: *Amphimedon queenslandica* (Porifera) as outgroup.
7. Porifera: *Mnemiopsis leidyi* (Ctenophora) as outgroup.
8. Brachiopoda: *Haliotis rufescens* (Mollusca) as outgroup.
9. Mollusca, Annelida, and Platyhelminthes: *Adineta vaga* (Rotifera) as outgroup.

```
# Create data frame of query species and outgroup
by_phylum <- function(df, taxon) {
    return(
        df |>
            dplyr::filter(phylum == taxon) |>
            dplyr::pull(species)
    )
}

arthropoda <- by_phylum(metadata_all$metazoa, "Arthropoda")
tardigrada <- by_phylum(metadata_all$metazoa, "Tardigrada")
```

```
nematoda <- by_phylum(metadata_all$metazoa, "Nematoda")
priapulida <- by_phylum(metadata_all$metazoa, "Priapulida")
xenacoelomorpha <- by_phylum(metadata_all$metazoa, "Xenacoelomorpha")
cnidaria <- by_phylum(metadata_all$metazoa, "Cnidaria")
placozoa <- by_phylum(metadata_all$metazoa, "Placozoa")
porifera <- by_phylum(metadata_all$metazoa, "Porifera")
brachiopoda <- by_phylum(metadata_all$metazoa, "Brachiopoda")
mollusca <- by_phylum(metadata_all$metazoa, "Mollusca")
echinodermata <- by_phylum(metadata_all$metazoa, "Echinodermata")
annelida <- by_phylum(metadata_all$metazoa, "Annelida")
platyhelminthes <- by_phylum(metadata_all$metazoa, "Platyhelminthes")
acanthocephala <- by_phylum(metadata_all$metazoa, "Acanthocephala")
chordata <- by_phylum(metadata_all$metazoa, "Chordata")
hemichordata <- by_phylum(metadata_all$metazoa, "Hemichordata")
rotifera <- by_phylum(metadata_all$metazoa, "Rotifera")

metazoa_outgroups <- metadata_all$metazoa |>
    filter(phylum != "Ctenophora") |>
    mutate(
        query = species,
        outgroup = case_when(
            species %in% arthropoda ~ "hypsibius_exemplaris_gca002082055v1",
            species %in% c(tardigrada, acanthocephala, rotifera) ~ "brugia_malayi",
            species %in% nematoda ~ "priapulus_caudatus_gca000485595v2",

            species %in% c(priapulida, echinodermata, chordata, hemichordata) ~
                "hofstenia_miamia",
            species %in% xenacoelomorpha ~ "actinia_tenebrosa_gca009602425v1",
            species %in% c(cnidaria, placozoa) ~
                "amphimedon_queenslandica_gca000090795v2rs",
            species %in% porifera ~ "mnemiopsis_leidyi",
            species %in% brachiopoda ~ "haliotis_rufescens_gca023055435v1rs",
            species %in% c(mollusca, annelida, platyhelminthes) ~ "adineta_vaga"
        )
    ) |>
    select(query, outgroup)
```

Identifying and classifying duplicates:

```
col_dir <- here("products", "result_files", "collinearity", "metazoa")
if(!dir.exists(col_dir)) { dir.create(col_dir, recursive = TRUE) }

# Identifying and classifying paralogs
metazoa_duplicates <- ensembl2duplicates(
    metadata = metadata_all$metazoa,
    ensembl = "metazoa",
    outgroups = metazoa_outgroups,
    collinearity_dir = col_dir,
    threads = 4
)
```

```r
# Classify genes into unique duplication modes
metazoa_duplicates_unique <- classify_genes(metazoa_duplicates)

# Save classification results
## Duplicate pairs
save(
    metazoa_duplicates,
    file = here("products", "result_files", "metazoa_duplicates.rda"),
    compress = "xz"
)

## Duplicated genes (unique duplication modes)
save(
    metazoa_duplicates_unique,
    file = here("products", "result_files", "metazoa_duplicates_unique.rda"),
    compress = "xz"
)
```

#### 3.5 Ensembl (Vertebrates)

Here, we will use the following outgroups per taxa:

1. Amniota: *Xenopus tropicalis* (Amphibia) as outgroup;
2. Amphibia: *Latimeria chalumnae* (West Indian Ocean coelacanth)
3. All bony and cartilaginous fish: *Eptatretus burgeri* (hagfish, Agnatha)
4. Agnatha: *Ciona intestinalis* (Tunicata)

```r
# Create a data frame of species and outgroups
amniota <- metadata_all$ensembl |>
    filter(
        class %in% c("Aves", "Mammalia", "Lepidosauria") |
            order %in% c("Testudines", "Crocodylia")
    ) |>
    pull(species)

amphibia <- metadata_all$ensembl |>
    filter(class == "Amphibia") |>
    pull(species)

fish <- metadata_all$ensembl |>
    filter(
        class %in% c("Actinopteri", "Chondrichthyes", "Cladistia") |
            order == "Coelacanthiformes"
    ) |>
    pull(species)

agnatha <- metadata_all$ensembl |>
    filter(
        class %in% c("Myxini", "Hyperoartia")
    ) |>
    pull(species)
```

```
ensembl_outgroups <- metadata_all$ensembl |>
    filter(!phylum %in% c("Nematoda", "Arthropoda", "Ascomycota")) |>
    mutate(
        query = species,
        outgroup = case_when(
            species %in% amniota ~ "xenopus_tropicalis",
            species %in% amphibia ~ "latimeria_chalumnae",
            species %in% fish ~ "eptatretus_burgeri",
            species %in% agnatha ~ "ciona_intestinalis"
        )
    ) |>
    select(query, outgroup) |>
    filter(!is.na(outgroup))
```

Identifying and classifying duplicates:

```
col_dir <- here("products", "result_files", "collinearity", "vertebrates")
if(!dir.exists(col_dir)) { dir.create(col_dir, recursive = TRUE) }

# Identifying and classifying paralogs
vertebrates_duplicates <- ensembl2duplicates(
    meta,
    ensembl = "ensembl",
    outgroups = ensembl_outgroups,
    collinearity_dir = col_dir,
    tsv_dir = "~/Documents/vertebrates_duplicates", # delete later
    threads = 4
)

# Classify genes into unique duplication modes
vertebrates_duplicates_unique <- classify_genes(vertebrates_duplicates)

# Save classification results
## Duplicate pairs
save(
    vertebrates_duplicates,
    file = here("products", "result_files", "vertebrates_duplicates.rda"),
    compress = "xz"
)

## Duplicated genes (unique duplication modes)
save(
    vertebrates_duplicates_unique,
    file = here("products", "result_files", "vertebrates_duplicates_unique.rda"),
    compress = "xz"
)
```

### Session info

This document was created under the following conditions:

```
## - Session info ---------------------------------------------------------------
##  setting  value
##  version  R version 4.3.2 (2023-10-31)
##  os       Ubuntu 22.04.3 LTS
##  system   x86_64, linux-gnu
##  ui       X11
##  language (EN)
##  collate  en_US.UTF-8
##  ctype    en_US.UTF-8
##  tz       Europe/Brussels
##  date     2024-02-27
##  pandoc   3.1.1 @ /usr/lib/rstudio/resources/app/bin/quarto/bin/tools/ (via rmarkdown)
##
## - Packages -------------------------------------------------------------------
##  package              * version     date (UTC) lib source
##  abind                  1.4-5       2016-07-21 [1] CRAN (R 4.3.2)
##  ade4                   1.7-22      2023-02-06 [1] CRAN (R 4.3.2)
##  AnnotationDbi          1.64.1      2023-11-03 [1] Bioconductor
##  ape                    5.7-1       2023-03-13 [1] CRAN (R 4.3.2)
##  Biobase                2.62.0      2023-10-24 [1] Bioconductor
##  BiocFileCache          2.10.1      2023-10-26 [1] Bioconductor
##  BiocGenerics           0.48.1      2023-11-01 [1] Bioconductor
##  BiocIO                 1.12.0      2023-10-24 [1] Bioconductor
##  BiocManager            1.30.22     2023-08-08 [1] CRAN (R 4.3.2)
##  BiocParallel           1.37.0      2024-01-19 [1] Github (Bioconductor/BiocParallel@79a1b2d)
##  BiocStyle            * 2.30.0      2023-10-24 [1] Bioconductor
##  biomaRt              * 2.58.2      2024-01-30 [1] Bioconductor 3.18 (R 4.3.2)
##  Biostrings             2.70.2      2024-01-28 [1] Bioconductor 3.18 (R 4.3.2)
##  bit                    4.0.5       2022-11-15 [1] CRAN (R 4.3.2)
##  bit64                  4.0.5       2020-08-30 [1] CRAN (R 4.3.2)
##  bitops                 1.0-7       2021-04-24 [1] CRAN (R 4.3.2)
##  blob                   1.2.4       2023-03-17 [1] CRAN (R 4.3.2)
##  bookdown               0.37        2023-12-01 [1] CRAN (R 4.3.2)
##  cachem                 1.0.8       2023-05-01 [1] CRAN (R 4.3.2)
##  cli                    3.6.2       2023-12-11 [1] CRAN (R 4.3.2)
##  coda                   0.19-4.1    2024-01-31 [1] CRAN (R 4.3.2)
##  codetools              0.2-19      2023-02-01 [4] CRAN (R 4.2.2)
##  colorspace             2.1-0       2023-01-23 [1] CRAN (R 4.3.2)
##  crayon                 1.5.2       2022-09-29 [1] CRAN (R 4.3.2)
##  curl                   5.2.0       2023-12-08 [1] CRAN (R 4.3.2)
##  DBI                    1.2.1       2024-01-12 [1] CRAN (R 4.3.2)
##  dbplyr                 2.4.0       2023-10-26 [1] CRAN (R 4.3.2)
##  DelayedArray           0.28.0      2023-10-24 [1] Bioconductor
##  digest                 0.6.34      2024-01-11 [1] CRAN (R 4.3.2)
##  doParallel             1.0.17      2022-02-07 [1] CRAN (R 4.3.2)
##  doubletrouble        * 1.3.4       2024-02-05 [1] Bioconductor
##  dplyr                * 1.1.4       2023-11-17 [1] CRAN (R 4.3.2)
```

```
## evaluate              0.23     2023-11-01 [1] CRAN (R 4.3.2)
## fansi                 1.0.6    2023-12-08 [1] CRAN (R 4.3.2)
## fastmap               1.1.1    2023-02-24 [1] CRAN (R 4.3.2)
## filelock              1.0.3    2023-12-11 [1] CRAN (R 4.3.2)
## forcats             * 1.0.0    2023-01-29 [1] CRAN (R 4.3.2)
## foreach               1.5.2    2022-02-02 [1] CRAN (R 4.3.2)
## generics              0.1.3    2022-07-05 [1] CRAN (R 4.3.2)
## GenomeInfoDb          1.38.6   2024-02-08 [1] Bioconductor 3.18 (R 4.3.2)
## GenomeInfoDbData      1.2.11   2023-12-21 [1] Bioconductor
## GenomicAlignments     1.38.2   2024-01-16 [1] Bioconductor 3.18 (R 4.3.2)
## GenomicFeatures       1.54.3   2024-01-31 [1] Bioconductor 3.18 (R 4.3.2)
## GenomicRanges         1.54.1   2023-10-29 [1] Bioconductor
## ggnetwork             0.5.13   2024-02-14 [1] CRAN (R 4.3.2)
## ggplot2             * 3.4.4    2023-10-12 [1] CRAN (R 4.3.2)
## glue                  1.7.0    2024-01-09 [1] CRAN (R 4.3.2)
## gtable                0.3.4    2023-08-21 [1] CRAN (R 4.3.2)
## here                * 1.0.1    2020-12-13 [1] CRAN (R 4.3.2)
## hms                   1.1.3    2023-03-21 [1] CRAN (R 4.3.2)
## htmltools             0.5.7    2023-11-03 [1] CRAN (R 4.3.2)
## htmlwidgets           1.6.4    2023-12-06 [1] CRAN (R 4.3.2)
## httr                  1.4.7    2023-08-15 [1] CRAN (R 4.3.2)
## igraph                2.0.1.1  2024-01-30 [1] CRAN (R 4.3.2)
## intergraph            2.0-4    2024-02-01 [1] CRAN (R 4.3.2)
## IRanges               2.36.0   2023-10-24 [1] Bioconductor
## iterators             1.0.14   2022-02-05 [1] CRAN (R 4.3.2)
## KEGGREST              1.42.0   2023-10-24 [1] Bioconductor
## knitr                 1.45     2023-10-30 [1] CRAN (R 4.3.2)
## lattice               0.22-5   2023-10-24 [4] CRAN (R 4.3.1)
## lifecycle             1.0.4    2023-11-07 [1] CRAN (R 4.3.2)
## lubridate           * 1.9.3    2023-09-27 [1] CRAN (R 4.3.2)
## magrittr              2.0.3    2022-03-30 [1] CRAN (R 4.3.2)
## MASS                  7.3-60   2023-05-04 [4] CRAN (R 4.3.1)
## Matrix                1.6-3    2023-11-14 [4] CRAN (R 4.3.2)
## MatrixGenerics        1.14.0   2023-10-24 [1] Bioconductor
## matrixStats           1.2.0    2023-12-11 [1] CRAN (R 4.3.2)
## mclust                6.0.1    2023-11-15 [1] CRAN (R 4.3.2)
## memoise               2.0.1    2021-11-26 [1] CRAN (R 4.3.2)
## MSA2dist              1.6.0    2023-10-24 [1] Bioconductor
## munsell               0.5.0    2018-06-12 [1] CRAN (R 4.3.2)
## network               1.18.2   2023-12-05 [1] CRAN (R 4.3.2)
## networkD3             0.4      2017-03-18 [1] CRAN (R 4.3.2)
## nlme                  3.1-163  2023-08-09 [4] CRAN (R 4.3.1)
## pheatmap              1.0.12   2019-01-04 [1] CRAN (R 4.3.2)
## pillar                1.9.0    2023-03-22 [1] CRAN (R 4.3.2)
## pkgconfig             2.0.3    2019-09-22 [1] CRAN (R 4.3.2)
## png                   0.1-8    2022-11-29 [1] CRAN (R 4.3.2)
## prettyunits           1.2.0    2023-09-24 [1] CRAN (R 4.3.2)
## progress              1.2.3    2023-12-06 [1] CRAN (R 4.3.2)
## purrr               * 1.0.2    2023-08-10 [1] CRAN (R 4.3.2)
## R6                    2.5.1    2021-08-19 [1] CRAN (R 4.3.2)
## rappdirs              0.3.3    2021-01-31 [1] CRAN (R 4.3.2)
```

```
##  RColorBrewer           1.1-3      2022-04-03 [1] CRAN (R 4.3.2)
##  Rcpp                   1.0.12     2024-01-09 [1] CRAN (R 4.3.2)
##  RCurl                  1.98-1.14  2024-01-09 [1] CRAN (R 4.3.2)
##  readr                * 2.1.5      2024-01-10 [1] CRAN (R 4.3.2)
##  restfulr               0.0.15     2022-06-16 [1] CRAN (R 4.3.2)
##  rjson                  0.2.21     2022-01-09 [1] CRAN (R 4.3.2)
##  rlang                  1.1.3      2024-01-10 [1] CRAN (R 4.3.2)
##  rmarkdown              2.25       2023-09-18 [1] CRAN (R 4.3.2)
##  rprojroot              2.0.4      2023-11-05 [1] CRAN (R 4.3.2)
##  Rsamtools              2.18.0     2023-10-24 [1] Bioconductor
##  RSQLite                2.3.5      2024-01-21 [1] CRAN (R 4.3.2)
##  rstudioapi             0.15.0     2023-07-07 [1] CRAN (R 4.3.2)
##  rtracklayer            1.62.0     2023-10-24 [1] Bioconductor
##  S4Arrays               1.2.0      2023-10-24 [1] Bioconductor
##  S4Vectors              0.40.2     2023-11-23 [1] Bioconductor 3.18 (R 4.3.2)
##  scales                 1.3.0      2023-11-28 [1] CRAN (R 4.3.2)
##  seqinr                 4.2-36     2023-12-08 [1] CRAN (R 4.3.2)
##  sessioninfo            1.2.2      2021-12-06 [1] CRAN (R 4.3.2)
##  SparseArray            1.2.4      2024-02-11 [1] Bioconductor 3.18 (R 4.3.2)
##  statnet.common         4.9.0      2023-05-24 [1] CRAN (R 4.3.2)
##  stringi                1.8.3      2023-12-11 [1] CRAN (R 4.3.2)
##  stringr              * 1.5.1      2023-11-14 [1] CRAN (R 4.3.2)
##  SummarizedExperiment   1.32.0     2023-10-24 [1] Bioconductor
##  syntenet             * 1.4.0      2023-10-24 [1] Bioconductor
##  tibble               * 3.2.1      2023-03-20 [1] CRAN (R 4.3.2)
##  tidyr                * 1.3.1      2024-01-24 [1] CRAN (R 4.3.2)
##  tidyselect             1.2.0      2022-10-10 [1] CRAN (R 4.3.2)
##  tidyverse            * 2.0.0      2023-02-22 [1] CRAN (R 4.3.2)
##  timechange             0.3.0      2024-01-18 [1] CRAN (R 4.3.2)
##  tzdb                   0.4.0      2023-05-12 [1] CRAN (R 4.3.2)
##  utf8                   1.2.4      2023-10-22 [1] CRAN (R 4.3.2)
##  vctrs                  0.6.5      2023-12-01 [1] CRAN (R 4.3.2)
##  withr                  3.0.0      2024-01-16 [1] CRAN (R 4.3.2)
##  xfun                   0.42       2024-02-08 [1] CRAN (R 4.3.2)
##  XML                    3.99-0.16.1 2024-01-22 [1] CRAN (R 4.3.2)
##  xml2                   1.3.6      2023-12-04 [1] CRAN (R 4.3.2)
##  XVector                0.42.0     2023-10-24 [1] Bioconductor
##  yaml                   2.3.8      2023-12-11 [1] CRAN (R 4.3.2)
##  zlibbioc               1.48.0     2023-10-24 [1] Bioconductor
##
##  [1] /home/faalm/R/x86_64-pc-linux-gnu-library/4.3
##  [2] /usr/local/lib/R/site-library
##  [3] /usr/lib/R/site-library
##  [4] /usr/lib/R/library
##
## ------------------------------------------------------------------------------
```

### Supplementary Text S3: Calculating substitution rates for selected Ensembl genomes

***Fabricio Almeida-Silva*** [1] ***and Yves Van de Peer*** [1]

[1]VIB-UGent Center for Plant Systems Biology, Ghent University, Ghent, Belgium

**27 February 2024**

**Supplementary Text S3: Calculating substitution rates for selected Ensembl genomes**

### 1 Introduction

Here, we will describe the code to calculate substitution rates for selected genomes in Ensembl and Ensembl Genomes instances using the Bioconductor package *doubletrouble*.

```
library(syntenet)
library(doubletrouble)
library(here)
## here() starts at /home/faalm/Dropbox/package_benchmarks/doubletrouble_paper
library(tidyverse)
## -- Attaching core tidyverse packages ------------------------ tidyverse 2.0.0 --
## v dplyr     1.1.4     v readr     2.1.5
## v forcats   1.0.0     v stringr   1.5.1
## v ggplot2   3.4.4     v tibble    3.2.1
## v lubridate 1.9.3     v tidyr     1.3.1
## v purrr     1.0.2
## -- Conflicts ------------------------------------------ tidyverse_conflicts() --
## x dplyr::filter() masks stats::filter()
## x dplyr::lag()    masks stats::lag()
## i Use the conflicted package (<http://conflicted.r-lib.org/>) to force all conflicts to become errors
library(BiocParallel)

set.seed(123) # for reproducibility
options(timeout = 1e10) # to allow download of big files

# Load helper functions
source(here("code", "utils.R"))
```

### 2 Data loading

Here, we will load the data frames of species metadata and the lists of duplicated gene pairs for each Ensembl instance.

```
# Load metadata
load(here("products", "result_files", "metadata_all.rda"))
names(metadata_all)
## [1] "fungi"    "plants"   "metazoa"  "protists" "ensembl"

# Load duplicates
load(here("products", "result_files", "fungi_duplicates.rda"))
load(here("products", "result_files", "plants_duplicates.rda"))
```

### 3 Calculating substitution rates

Next, we will calculate substitution rates ($K_a$, $K_s$, and $K_a/K_s$) for duplicate pairs in all selected species, namely:

1. Three fungi species (*Saccharomyces cerevisiae*, *Candida glabrata*, and *Schizosaccharomyces pombe*).

2. Four legume species (*Glycine max*, *Phaseolus vulgaris*, *Vitis vinifera*, *Selaginella moellendorffii*).

```
# Fungi - S. cerevisiae, Candida glabrata, and Schizosaccharomyces pombe
## Download CDS
selected_fungi <- c(
    "saccharomyces_cerevisiae", "candida_glabrata", "schizosaccharomyces_pombe"
)
fungi_cds <- get_cds_ensembl(selected_fungi, ensembl = "fungi")

## Calculate substitution rates
fungi_kaks <- pairs2kaks(
    gene_pairs_list = fungi_duplicates[selected_fungi],
    cds = fungi_cds,
    bp_param = BiocParallel::SnowParam(workers = 8)
)

# Plants - Glycine max, Phaseolus vulgaris, Vitis vinifera, and Selaginella moellendorffii
## Download CDS
selected_plants <- c(
    "glycine_max", "phaseolus_vulgaris", "vitis_vinifera",
    "selaginella_moellendorffii"
)

plants_cds <- get_cds_ensembl(selected_plants, ensembl = "plants")

## Calculate substitution rates
plants_duplicates <- plants_duplicates[selected_plants]

plants_kaks <- pairs2kaks(
    gene_pairs_list = plants_duplicates,
    cds = plants_cds,
    bp_param = BiocParallel::SnowParam(workers = 8)
)
```

Saving objects as .rda files:

```
save(
    fungi_kaks, compress = "xz",
    file = here("products", "result_files", "fungi_kaks.rda")
)

save(
    plants_kaks, compress = "xz",
    file = here("products", "result_files", "plants_kaks.rda")
)
```

### Session info

This document was created under the following conditions:

```
## - Session info ---------------------------------------------------------------
##  setting  value
##  version  R version 4.3.2 (2023-10-31)
##  os       Ubuntu 22.04.3 LTS
##  system   x86_64, linux-gnu
##  ui       X11
##  language (EN)
##  collate  en_US.UTF-8
##  ctype    en_US.UTF-8
##  tz       Europe/Brussels
##  date     2024-02-27
##  pandoc   3.1.1 @ /usr/lib/rstudio/resources/app/bin/quarto/bin/tools/ (via rmarkdown)
##
## - Packages -------------------------------------------------------------------
##  package              * version     date (UTC) lib source
##  abind                  1.4-5       2016-07-21 [1] CRAN (R 4.3.2)
##  ade4                   1.7-22      2023-02-06 [1] CRAN (R 4.3.2)
##  AnnotationDbi          1.64.1      2023-11-03 [1] Bioconductor
##  ape                    5.7-1       2023-03-13 [1] CRAN (R 4.3.2)
##  Biobase                2.62.0      2023-10-24 [1] Bioconductor
##  BiocFileCache          2.10.1      2023-10-26 [1] Bioconductor
##  BiocGenerics           0.48.1      2023-11-01 [1] Bioconductor
##  BiocIO                 1.12.0      2023-10-24 [1] Bioconductor
##  BiocManager            1.30.22     2023-08-08 [1] CRAN (R 4.3.2)
##  BiocParallel         * 1.37.0      2024-01-19 [1] Github (Bioconductor/BiocParallel@79a1b2d)
##  BiocStyle            * 2.30.0      2023-10-24 [1] Bioconductor
##  biomaRt                2.58.2      2024-01-30 [1] Bioconductor 3.18 (R 4.3.2)
##  Biostrings             2.70.2      2024-01-28 [1] Bioconductor 3.18 (R 4.3.2)
##  bit                    4.0.5       2022-11-15 [1] CRAN (R 4.3.2)
##  bit64                  4.0.5       2020-08-30 [1] CRAN (R 4.3.2)
##  bitops                 1.0-7       2021-04-24 [1] CRAN (R 4.3.2)
##  blob                   1.2.4       2023-03-17 [1] CRAN (R 4.3.2)
##  bookdown               0.37        2023-12-01 [1] CRAN (R 4.3.2)
##  cachem                 1.0.8       2023-05-01 [1] CRAN (R 4.3.2)
##  cli                    3.6.2       2023-12-11 [1] CRAN (R 4.3.2)
##  coda                   0.19-4.1    2024-01-31 [1] CRAN (R 4.3.2)
##  codetools              0.2-19      2023-02-01 [4] CRAN (R 4.2.2)
##  colorspace             2.1-0       2023-01-23 [1] CRAN (R 4.3.2)
##  crayon                 1.5.2       2022-09-29 [1] CRAN (R 4.3.2)
##  curl                   5.2.0       2023-12-08 [1] CRAN (R 4.3.2)
##  DBI                    1.2.1       2024-01-12 [1] CRAN (R 4.3.2)
##  dbplyr                 2.4.0       2023-10-26 [1] CRAN (R 4.3.2)
##  DelayedArray           0.28.0      2023-10-24 [1] Bioconductor
##  digest                 0.6.34      2024-01-11 [1] CRAN (R 4.3.2)
##  doParallel             1.0.17      2022-02-07 [1] CRAN (R 4.3.2)
##  doubletrouble        * 1.3.4       2024-02-05 [1] Bioconductor
##  dplyr                * 1.1.4       2023-11-17 [1] CRAN (R 4.3.2)
##  evaluate               0.23        2023-11-01 [1] CRAN (R 4.3.2)
##  fansi                  1.0.6       2023-12-08 [1] CRAN (R 4.3.2)
##  fastmap                1.1.1       2023-02-24 [1] CRAN (R 4.3.2)
##  filelock               1.0.3       2023-12-11 [1] CRAN (R 4.3.2)
```

```
## forcats            * 1.0.0     2023-01-29 [1] CRAN (R 4.3.2)
## foreach              1.5.2     2022-02-02 [1] CRAN (R 4.3.2)
## generics             0.1.3     2022-07-05 [1] CRAN (R 4.3.2)
## GenomeInfoDb         1.38.6    2024-02-08 [1] Bioconductor 3.18 (R 4.3.2)
## GenomeInfoDbData     1.2.11    2023-12-21 [1] Bioconductor
## GenomicAlignments    1.38.2    2024-01-16 [1] Bioconductor 3.18 (R 4.3.2)
## GenomicFeatures      1.54.3    2024-01-31 [1] Bioconductor 3.18 (R 4.3.2)
## GenomicRanges        1.54.1    2023-10-29 [1] Bioconductor
## ggnetwork            0.5.13    2024-02-14 [1] CRAN (R 4.3.2)
## ggplot2            * 3.4.4     2023-10-12 [1] CRAN (R 4.3.2)
## glue                 1.7.0     2024-01-09 [1] CRAN (R 4.3.2)
## gtable               0.3.4     2023-08-21 [1] CRAN (R 4.3.2)
## here               * 1.0.1     2020-12-13 [1] CRAN (R 4.3.2)
## hms                  1.1.3     2023-03-21 [1] CRAN (R 4.3.2)
## htmltools            0.5.7     2023-11-03 [1] CRAN (R 4.3.2)
## htmlwidgets          1.6.4     2023-12-06 [1] CRAN (R 4.3.2)
## httr                 1.4.7     2023-08-15 [1] CRAN (R 4.3.2)
## igraph               2.0.1.1   2024-01-30 [1] CRAN (R 4.3.2)
## intergraph           2.0-4     2024-02-01 [1] CRAN (R 4.3.2)
## IRanges              2.36.0    2023-10-24 [1] Bioconductor
## iterators            1.0.14    2022-02-05 [1] CRAN (R 4.3.2)
## KEGGREST             1.42.0    2023-10-24 [1] Bioconductor
## knitr                1.45      2023-10-30 [1] CRAN (R 4.3.2)
## lattice              0.22-5    2023-10-24 [4] CRAN (R 4.3.1)
## lifecycle            1.0.4     2023-11-07 [1] CRAN (R 4.3.2)
## lubridate          * 1.9.3     2023-09-27 [1] CRAN (R 4.3.2)
## magrittr             2.0.3     2022-03-30 [1] CRAN (R 4.3.2)
## MASS                 7.3-60    2023-05-04 [4] CRAN (R 4.3.1)
## Matrix               1.6-3     2023-11-14 [4] CRAN (R 4.3.2)
## MatrixGenerics       1.14.0    2023-10-24 [1] Bioconductor
## matrixStats          1.2.0     2023-12-11 [1] CRAN (R 4.3.2)
## mclust               6.0.1     2023-11-15 [1] CRAN (R 4.3.2)
## memoise              2.0.1     2021-11-26 [1] CRAN (R 4.3.2)
## MSA2dist             1.6.0     2023-10-24 [1] Bioconductor
## munsell              0.5.0     2018-06-12 [1] CRAN (R 4.3.2)
## network              1.18.2    2023-12-05 [1] CRAN (R 4.3.2)
## networkD3            0.4       2017-03-18 [1] CRAN (R 4.3.2)
## nlme                 3.1-163   2023-08-09 [4] CRAN (R 4.3.1)
## pheatmap             1.0.12    2019-01-04 [1] CRAN (R 4.3.2)
## pillar               1.9.0     2023-03-22 [1] CRAN (R 4.3.2)
## pkgconfig            2.0.3     2019-09-22 [1] CRAN (R 4.3.2)
## png                  0.1-8     2022-11-29 [1] CRAN (R 4.3.2)
## prettyunits          1.2.0     2023-09-24 [1] CRAN (R 4.3.2)
## progress             1.2.3     2023-12-06 [1] CRAN (R 4.3.2)
## purrr              * 1.0.2     2023-08-10 [1] CRAN (R 4.3.2)
## R6                   2.5.1     2021-08-19 [1] CRAN (R 4.3.2)
## rappdirs             0.3.3     2021-01-31 [1] CRAN (R 4.3.2)
## RColorBrewer         1.1-3     2022-04-03 [1] CRAN (R 4.3.2)
## Rcpp                 1.0.12    2024-01-09 [1] CRAN (R 4.3.2)
## RCurl                1.98-1.14 2024-01-09 [1] CRAN (R 4.3.2)
## readr              * 2.1.5     2024-01-10 [1] CRAN (R 4.3.2)
```

```
## restfulr               0.0.15      2022-06-16 [1] CRAN (R 4.3.2)
## rjson                  0.2.21      2022-01-09 [1] CRAN (R 4.3.2)
## rlang                  1.1.3       2024-01-10 [1] CRAN (R 4.3.2)
## rmarkdown              2.25        2023-09-18 [1] CRAN (R 4.3.2)
## rprojroot              2.0.4       2023-11-05 [1] CRAN (R 4.3.2)
## Rsamtools              2.18.0      2023-10-24 [1] Bioconductor
## RSQLite                2.3.5       2024-01-21 [1] CRAN (R 4.3.2)
## rstudioapi             0.15.0      2023-07-07 [1] CRAN (R 4.3.2)
## rtracklayer            1.62.0      2023-10-24 [1] Bioconductor
## S4Arrays               1.2.0       2023-10-24 [1] Bioconductor
## S4Vectors              0.40.2      2023-11-23 [1] Bioconductor 3.18 (R 4.3.2)
## scales                 1.3.0       2023-11-28 [1] CRAN (R 4.3.2)
## seqinr                 4.2-36      2023-12-08 [1] CRAN (R 4.3.2)
## sessioninfo            1.2.2       2021-12-06 [1] CRAN (R 4.3.2)
## SparseArray            1.2.4       2024-02-11 [1] Bioconductor 3.18 (R 4.3.2)
## statnet.common         4.9.0       2023-05-24 [1] CRAN (R 4.3.2)
## stringi                1.8.3       2023-12-11 [1] CRAN (R 4.3.2)
## stringr              * 1.5.1       2023-11-14 [1] CRAN (R 4.3.2)
## SummarizedExperiment   1.32.0      2023-10-24 [1] Bioconductor
## syntenet             * 1.4.0       2023-10-24 [1] Bioconductor
## tibble               * 3.2.1       2023-03-20 [1] CRAN (R 4.3.2)
## tidyr                * 1.3.1       2024-01-24 [1] CRAN (R 4.3.2)
## tidyselect             1.2.0       2022-10-10 [1] CRAN (R 4.3.2)
## tidyverse            * 2.0.0       2023-02-22 [1] CRAN (R 4.3.2)
## timechange             0.3.0       2024-01-18 [1] CRAN (R 4.3.2)
## tzdb                   0.4.0       2023-05-12 [1] CRAN (R 4.3.2)
## utf8                   1.2.4       2023-10-22 [1] CRAN (R 4.3.2)
## vctrs                  0.6.5       2023-12-01 [1] CRAN (R 4.3.2)
## withr                  3.0.0       2024-01-16 [1] CRAN (R 4.3.2)
## xfun                   0.42        2024-02-08 [1] CRAN (R 4.3.2)
## XML                    3.99-0.16.1 2024-01-22 [1] CRAN (R 4.3.2)
## xml2                   1.3.6       2023-12-04 [1] CRAN (R 4.3.2)
## XVector                0.42.0      2023-10-24 [1] Bioconductor
## yaml                   2.3.8       2023-12-11 [1] CRAN (R 4.3.2)
## zlibbioc               1.48.0      2023-10-24 [1] Bioconductor
##
## [1] /home/faalm/R/x86_64-pc-linux-gnu-library/4.3
## [2] /usr/local/lib/R/site-library
## [3] /usr/lib/R/site-library
## [4] /usr/lib/R/library
##
## ------------------------------------------------------------------------------
```

### Supplementary Text S4: Visual exploration of duplicated genes across the Eukarya tree of life

***Fabricio Almeida-Silva*[1] *and Yves Van de Peer*[1]**

[1]VIB-UGent Center for Plant Systems Biology, Ghent University, Ghent, Belgium

**27 February 2024**

### 1 Introduction

Here, we will describe the code to perform exploratory data analyses on the duplicated gene frequencies in genomes from Ensembl instances.

To start, let's load the required data and packages.

```
set.seed(123) # for reproducibility

# Load required packages
library(doubletrouble)
## Registered S3 method overwritten by 'ggnetwork':
##   method         from
##   fortify.igraph ggtree
library(here)
## here() starts at /home/faalm/Dropbox/package_benchmarks/doubletrouble_paper
library(ggtree)
## ggtree v3.10.0 For help: https://yulab-smu.top/treedata-book/
##
## If you use the ggtree package suite in published research, please cite
## the appropriate paper(s):
##
## Guangchuang Yu, David Smith, Huachen Zhu, Yi Guan, Tommy Tsan-Yuk Lam.
## ggtree: an R package for visualization and annotation of phylogenetic
## trees with their covariates and other associated data. Methods in
## Ecology and Evolution. 2017, 8(1):28-36. doi:10.1111/2041-210X.12628
##
## G Yu. Data Integration, Manipulation and Visualization of Phylogenetic
## Trees (1st ed.). Chapman and Hall/CRC. 2022. ISBN: 9781032233574
##
## Shuangbin Xu, Lin Li, Xiao Luo, Meijun Chen, Wenli Tang, Li Zhan, Zehan
## Dai, Tommy T. Lam, Yi Guan, Guangchuang Yu. Ggtree: A serialized data
## object for visualization of a phylogenetic tree and annotation data.
## iMeta 2022, 1(4):e56. doi:10.1002/imt2.56
library(tidyverse)
## -- Attaching core tidyverse packages ------------------------ tidyverse 2.0.0 --
## v dplyr     1.1.4     v readr     2.1.5
## v forcats   1.0.0     v stringr   1.5.1
## v ggplot2   3.4.4     v tibble    3.2.1
## v lubridate 1.9.3     v tidyr     1.3.1
## v purrr     1.0.2
## -- Conflicts ------------------------------------------ tidyverse_conflicts() --
## x tidyr::expand() masks ggtree::expand()
## x dplyr::filter() masks stats::filter()
## x dplyr::lag()    masks stats::lag()
## i Use the conflicted package (<http://conflicted.r-lib.org/>) to force all conflicts to become errors
library(patchwork)

source(here("code", "utils.R"))
source(here("code", "utils_visualization.R"))
```

#### 2 Loading data

First, we will load objects with species trees, duplicates per species, and BUSCO scores.

```
# Load metadata
load(here("products", "result_files", "metadata_all.rda"))

# Load BUSCO scores
load(here("products", "result_files", "busco_scores", "fungi_busco_scores.rda"))
load(here("products", "result_files", "busco_scores", "protists_busco_scores.rda"))
load(here("products", "result_files", "busco_scores", "plants_busco_scores.rda"))
load(here("products", "result_files", "busco_scores", "metazoa_busco_scores.rda"))
load(here("products", "result_files", "busco_scores", "vertebrates_busco_scores.rda"))

# Load trees
load(here("products", "result_files", "trees", "fungi_busco_trees.rda"))
load(here("products", "result_files", "trees", "protists_busco_trees.rda"))
load(here("products", "result_files", "trees", "plants_busco_trees.rda"))
load(here("products", "result_files", "trees", "metazoa_busco_trees.rda"))
load(here("products", "result_files", "trees", "vertebrates_busco_trees.rda"))

# Load duplicated genes
load(here("products", "result_files", "fungi_duplicates_unique.rda"))
load(here("products", "result_files", "protists_duplicates_unique.rda"))
load(here("products", "result_files", "plants_duplicates_unique.rda"))
load(here("products", "result_files", "vertebrates_duplicates_unique.rda"))
load(here("products", "result_files", "metazoa_duplicates_unique.rda"))

# Load substitution rates for plants
load(here("products", "result_files", "plants_kaks.rda"))
```

#### 3 Visualizing the frequency of duplicated genes by mode

Now, we will visualize the frequency of duplicated genes by mode for each species. For that, we will first convert the list of duplicates into a long-formatted data frame, and clean tip labels in our species trees.

```
# Rename tip labels of trees
tree_fungi <- fungi_busco_trees$conc
tree_fungi$tip.label <- gsub("\\.", "_", tree_fungi$tip.label)

tree_protists <- protists_busco_trees$conc
tree_protists$tip.label <- gsub("\\.", "_", tree_protists$tip.label)

tree_plants <- plants_busco_trees$conc
tree_plants$tip.label <- gsub("\\.", "_", tree_plants$tip.label)

tree_metazoa <- metazoa_busco_trees$conc
```

```
tree_metazoa$tip.label <- gsub("\\.", "_", tree_metazoa$tip.label)

tree_vertebrates <- vertebrates_busco_trees$conc
tree_vertebrates$tip.label <- gsub("\\.", "_", tree_vertebrates$tip.label)

# Get count tables
counts_fungi <- duplicates2counts(fungi_duplicates_unique)
counts_protists <- duplicates2counts(protists_duplicates_unique)
counts_plants <- duplicates2counts(plants_duplicates_unique)
counts_vertebrates <- duplicates2counts(vertebrates_duplicates_unique)
counts_metazoa <- duplicates2counts(metazoa_duplicates_unique)
```

Now, we will plot the trees with data for each Ensembl instance.

```
# Fungi
p_fungi_tree <- plot_tree_taxa(
    tree = tree_fungi,
    metadata = metadata_all$fungi,
    taxon = "phylum",
    text_size = 2.5
)

p_fungi <- wrap_plots(
    # Plot 1: Species tree
    p_fungi_tree,
    # Plot 2: Duplicate relative frequency by mode
    plot_duplicate_freqs(
        counts_fungi |>
            mutate(
                species = factor(species, levels = rev(get_taxa_name(p_fungi_tree)))
            ),
        plot_type = "stack_percent"
    ) +
        theme(
            axis.text.y = element_blank(),
            axis.ticks.y = element_blank()
        ) +
        labs(y = NULL),
    widths = c(1, 4)
) +
    plot_annotation(title = "Fungi") &
    theme(plot.margin = margin(2, 0, 0, 2))

# Protists
p_protists_tree <- plot_tree_taxa(
    tree = tree_protists,
    metadata = metadata_all$protists |>
        filter(phylum != "Evosea"),
    taxon = "phylum",
    min_n_lab = 2,
    padding_text = 0.2,
```

```
        text_size = 2.5
    )
    p_protists <- wrap_plots(
        # Plot 1: Species tree
        p_protists_tree,
        # Plot 2: Duplicate relative frequency by mode
        plot_duplicate_freqs(
            counts_protists |>
                mutate(
                    species = factor(species, levels = rev(get_taxa_name(p_protists_tree)))
                ),
            plot_type = "stack_percent") +
            theme(
                axis.text.y = element_blank(),
                axis.ticks.y = element_blank()
            ) +
            labs(y = NULL),
        widths = c(1, 4)
    ) +
        plot_annotation(title = "Protists") &
        theme(plot.margin = margin(2, 0, 0, 2))

    # Plants
    p_plants_tree <- plot_tree_taxa(
        tree = tree_plants,
        metadata = metadata_all$plants,
        taxon = "order",
        min_n_lab = 3,
        text_size = 2.5
    )
    p_plants <- wrap_plots(
        # Plot 1: Species tree
        p_plants_tree,
        # Plot 2: Duplicate relative frequency by mode
        plot_duplicate_freqs(
            counts_plants |>
                mutate(
                    species = factor(species, levels = rev(get_taxa_name(p_plants_tree)))
                ),
            plot_type = "stack_percent") +
            theme(
                axis.text.y = element_blank(),
                axis.ticks.y = element_blank()
            ) +
            labs(y = NULL),
        widths = c(1, 4)
    ) +
        plot_annotation(title = "Plants") &
        theme(plot.margin = margin(2, 0, 0, 2))
```

```
    # Metazoa
    p_metazoa_tree <- plot_tree_taxa(
        tree = tree_metazoa,
        metadata = metadata_all$metazoa |>
            filter(class != "Myxozoa"),
        taxon = "phylum",
        min_n = 2,
        text_size = 2.2,
        padding_text = 2
    )
    p_metazoa <- wrap_plots(
        # Plot 1: Species tree
        p_metazoa_tree,
        # Plot 2: Duplicate relative frequency by mode
        plot_duplicate_freqs(
            counts_metazoa |>
                mutate(
                    species = factor(species, levels = rev(get_taxa_name(p_metazoa_tree)))
                ),
            plot_type = "stack_percent") +
            theme(
                axis.text.y = element_blank(),
                axis.ticks.y = element_blank()
            ) +
            labs(y = NULL),
        widths = c(1, 4)
    ) +
        plot_annotation(title = "Metazoa") &
        theme(plot.margin = margin(2, 0, 0, 2))

    # Vertebrates
    p_vertebrates_tree <- plot_tree_taxa(
        tree = tree_vertebrates,
        metadata = metadata_all$ensembl |>
            mutate(class = replace_na(class, "Other")),
        taxon = "class",
        min_n = 2,
        text_size = 2.5
    )
    p_vertebrates <- wrap_plots(
        # Plot 1: Species tree
        p_vertebrates_tree,
        # Plot 2: Duplicate relative frequency by mode
        plot_duplicate_freqs(
            counts_vertebrates |>
                mutate(
                    species = factor(species, levels = rev(get_taxa_name(p_vertebrates_tree)))
                ),
            plot_type = "stack_percent") +
            theme(
                axis.text.y = element_blank(),
```

```
                axis.ticks.y = element_blank()
            ) +
            labs(y = NULL),
        widths = c(1, 4)
    ) +
        plot_annotation(title = "Vertebrates") &
        theme(plot.margin = margin(2, 0, 0, 2))

    # Combining all figures into one
    p_duplicates_all_ensembl <- wrap_plots(
        wrap_plots(
            p_protists +
                theme(legend.position = "none") +
                labs(title = "Protists", x = NULL),
            p_fungi +
                theme(legend.position = "none") +
                ggtitle("Fungi"),
            nrow = 2, heights = c(1, 2)
        ),
        p_plants + theme(legend.position = "none") + ggtitle("Plants"),
        p_metazoa + theme(legend.position = "none") + ggtitle("Metazoa (Invertebrates)"),
        p_vertebrates + ggtitle("Ensembl (Vertebrates)"),
        nrow = 1
    ) +
        plot_layout(axis_titles = "collect")
```

```
    p_duplicates_all_ensembl
```

#### Supplementary Text S4: Visual exploration of duplicated genes across the Eukarya tree of life

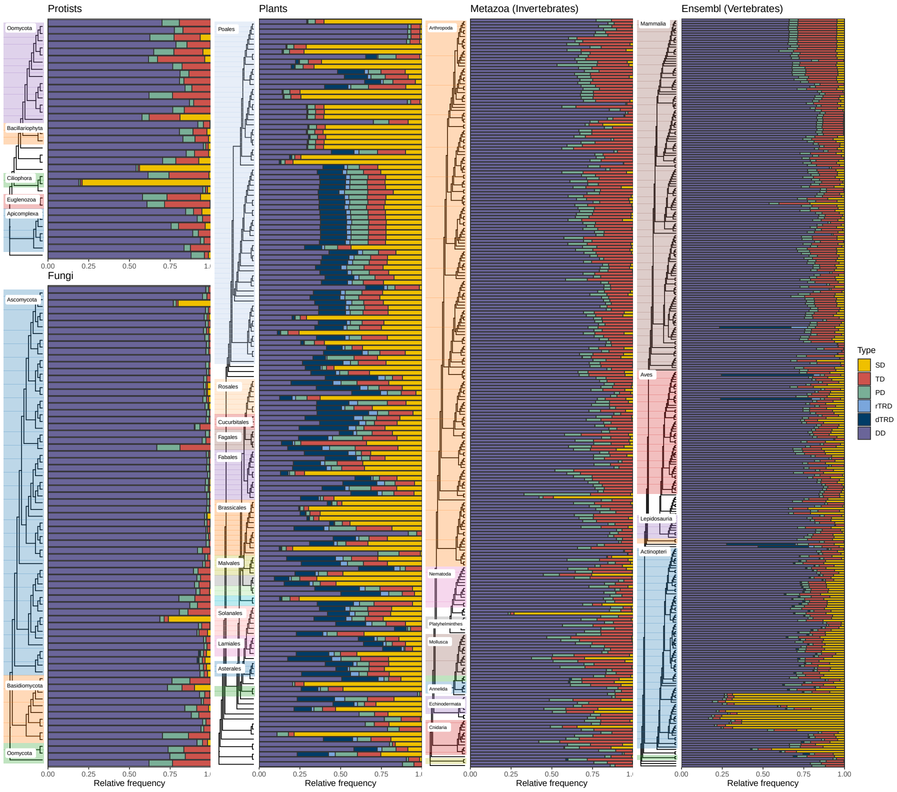

By visually comparing the Ensembl instances, we can see that plant genomes have a much greater abundance of segmental duplicates, possibly due to pervasive whole-genome duplication events. However, other major branches of the Eukarya tree of life also have particular species with a high proportion of SD-derived genes. Notably, while SD events are widespread in plants, vertebrate species with high proportions of SD-derived genes are concentrated in a particular branch (teleost fishes). To investigate that, we will highlight species for which at least 20% of the duplicated genes derived from segmental duplications.

```
# For each Ensembl instance, show species with >=20% of genes derived from SD
## Define helper function
sd_abundant <- function(count_table, min = 20) {

    perc_table <- count_table |>
        group_by(species) |>
        mutate(percentage = (n / sum(n)) * 100) |>
        ungroup() |>
        filter(type == "SD", percentage >= min)

    return(perc_table)
}

# Get a table of SD-abundant species for each instance
sd_abundant_spp <- bind_rows(
    sd_abundant(counts_fungi) |> mutate(instance = "fungi"),
    sd_abundant(counts_protists) |> mutate(instance = "protists"),
    sd_abundant(counts_plants) |> mutate(instance = "plants"),
```

```
    sd_abundant(counts_vertebrates) |> mutate(instance = "vertebrates"),
    sd_abundant(counts_metazoa) |> mutate(instance = "metazoa")
) |>
    as.data.frame()
```

Then, let's summarize the frequencies (absolute and relative) in a table.

```
# How many species per instance?
sd_abundant_spp |>
    count(instance) |>
    mutate(
        percentage = n / c(
            nrow(metadata_all$fungi),
            nrow(metadata_all$metazoa),
            nrow(metadata_all$plants),
            nrow(metadata_all$protists),
            nrow(metadata_all$ensembl)
        ) * 100
    )
##      instance  n percentage
## 1       fungi  2   2.857143
## 2     metazoa  7   2.766798
## 3      plants 94  63.087248
## 4    protists  2   6.060606
## 5 vertebrates 21   6.624606
```

Once again, our findings highlight the abundance of large-scale duplications in plant genomes, as segmental duplications contributed to 20% of the duplicated genes in 94 species (63%). Next, let's print all SD-abundant species.

```
# Show all species
knitr::kable(sd_abundant_spp)
```

| type | n | species | percentage | instance |
|---|---|---|---|---|
| SD | 2138 | fusarium_oxysporum | 20.16030 | fungi |
| SD | 683 | saccharomyces_cerevisiae | 26.16858 | fungi |
| SD | 14351 | emiliania_huxleyi | 43.78509 | protists |
| SD | 27661 | paramecium_tetraurelia | 79.03369 | protists |
| SD | 22430 | actinidia_chinensis | 72.54908 | plants |
| SD | 4357 | ananas_comosus | 22.15048 | plants |
| SD | 5885 | arabidopsis_halleri | 21.37202 | plants |
| SD | 7530 | arabidopsis_thaliana | 33.29796 | plants |
| SD | 42997 | avena_sativa_ot3098 | 74.43306 | plants |
| SD | 63492 | avena_sativa_sang | 79.22833 | plants |
| SD | 7461 | brachypodium_distachyon | 27.24285 | plants |
| SD | 48381 | brassica_juncea | 69.66407 | plants |
| SD | 59918 | brassica_napus | 63.31417 | plants |
| SD | 24314 | brassica_oleracea | 44.30232 | plants |
| SD | 23541 | brassica_rapa | 62.58907 | plants |
| SD | 19282 | brassica_rapa_ro18 | 49.29567 | plants |
| SD | 69355 | camelina_sativa | 79.45628 | plants |
| SD | 15785 | chenopodium_quinoa | 48.57371 | plants |
| SD | 3600 | citrullus_lanatus | 22.76608 | plants |
| SD | 4084 | coffea_canephora | 20.60545 | plants |
| SD | 3317 | cynara_cardunculus | 20.20959 | plants |
| SD | 41246 | digitaria_exilis | 75.51999 | plants |
| SD | 8809 | dioscorea_rotundata | 32.30764 | plants |
| SD | 66262 | echinochloa_crusgalli | 70.97016 | plants |
| SD | 13447 | eragrostis_curvula | 27.40650 | plants |
| SD | 46488 | eucalyptus_grandis | 75.45773 | plants |
| SD | 6846 | ficus_carica | 30.61580 | plants |
| SD | 6747 | galdieria_sulphuraria | 25.40573 | plants |
| SD | 36994 | glycine_max | 72.19468 | plants |
| SD | 16290 | gossypium_raimondii | 48.65882 | plants |
| SD | 13432 | helianthus_annuus | 23.21425 | plants |
| SD | 9537 | ipomoea_triloba | 35.65100 | plants |
| SD | 13096 | juglans_regia | 34.16556 | plants |
| SD | 7374 | kalanchoe_fedtschenkoi | 28.37900 | plants |
| SD | 6757 | lactuca_sativa | 20.52178 | plants |
| SD | 5758 | leersia_perrieri | 25.97321 | plants |
| SD | 16167 | lupinus_angustifolius | 54.14448 | plants |
| SD | 22987 | malus_domestica_golden | 62.58031 | plants |
| SD | 14278 | manihot_esculenta | 52.81693 | plants |
| SD | 16170 | musa_acuminata | 54.10017 | plants |
| SD | 4586 | nymphaea_colorata | 20.98472 | plants |
| SD | 5092 | oryza_barthii | 20.55464 | plants |
| SD | 5078 | oryza_brachyantha | 23.83366 | plants |
| SD | 6054 | oryza_glaberrima | 23.77567 | plants |
| SD | 5454 | oryza_glumipatula | 21.46484 | plants |
| SD | 5847 | oryza_punctata | 24.25437 | plants |
| SD | 5587 | oryza_rufipogon | 21.24335 | plants |
| SD | 6096 | oryza_sativa | 23.94344 | plants |
| SD | 6136 | oryza_sativa_arc | 22.25769 | plants |
| SD | 6139 | oryza_sativa_azucena | 22.19370 | plants |
| SD | 6186 | oryza_sativa_chaomeo | 22.05583 | plants |
| SD | 6125 | oryza_sativa_gobolsailbalam | 22.38506 | plants |
| SD | 6207 | oryza_sativa_ir64 | 22.73876 | plants |
| SD | 6131 | oryza_sativa_ketannangka | 22.16238 | plants |
| SD | 6479 | oryza_sativa_khaoyaiguang | 23.36964 | plants |
| SD | 6144 | oryza_sativa_larhamugad | 22.46107 | plants |

### 4 BUSCO scores

Next, we will test whether the percentage of segmental duplicates in genomes is associated with the percentage of complete BUSCOs. In other words, we want to find out whether the low percentages of SD gene pairs is due to genome fragmentation.

```
# Define function to plot association between % SD and % complete BUSCOs
plot_busco_sd_assoc <- function(busco_df, counts_table) {

    p <- busco_df |>
        filter(Class %in% c("Complete_SC", "Complete_duplicate")) |>
        mutate(species = str_replace_all(File, "\\.fa", "")) |>
        mutate(species = str_replace_all(species, "\\.", "_")) |>
        group_by(species) |>
        summarise(complete_BUSCOs = sum(Frequency)) |>
        inner_join(sd_abundant(counts_table, min = 0)) |>
        ggpubr::ggscatter(
            x = "complete_BUSCOs", y = "percentage",
            color = "deepskyblue4", alpha = 0.4,
            add = "reg.line", add.params = list(
                color = "black", fill = "lightgray"
            ),
            conf.int = TRUE,
            cor.coef = TRUE,
            cor.coeff.args = list(
                method = "pearson", label.x = 3, label.sep = "\n"
            )
        ) +
        labs(x = "% complete BUSCOs", y = "% SD duplicates")

    return(p)
}

# Fungi
p_busco_association <- patchwork::wrap_plots(
    plot_busco_sd_assoc(fungi_busco_scores, counts_fungi) +
        labs(title = "Fungi"),
    plot_busco_sd_assoc(plants_busco_scores, counts_plants) +
        labs(title = "Plants"),
    plot_busco_sd_assoc(protists_busco_scores, counts_protists) +
        labs(title = "Protists"),
    plot_busco_sd_assoc(metazoa_busco_scores, counts_metazoa) +
        labs(title = "Metazoa"),
    plot_busco_sd_assoc(vertebrates_busco_scores, counts_vertebrates) +
        labs(title = "Vertebrates"),
    nrow = 1
)
```

```
p_busco_association
```

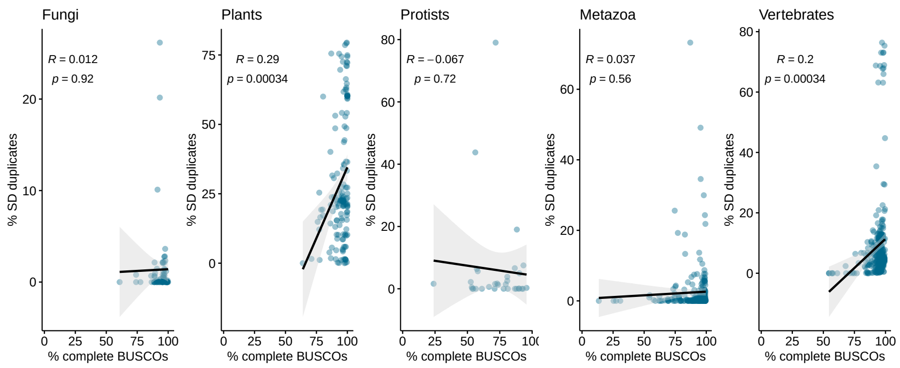

There is weak or no association between the percentage of complete BUSCOs and the percentage of SD-derived genes.

### 5 Visualizing substitution rates for selected plant species

Here, we will first visualize $K_s$ distributions for *Glycine max* and *Phaseolus vulgaris* by mode of duplication.

```
# G. max
gmax_ks_distro <- plot_ks_distro(
    plants_kaks$glycine_max, max_ks = 2, bytype = TRUE, binwidth = 0.03
) +
    labs(title = NULL, y = NULL)

# P. vulgaris
pvu_ks_distro <- plot_ks_distro(
    plants_kaks$phaseolus_vulgaris, max_ks = 2, bytype = TRUE, binwidth = 0.03
) +
    labs(title = NULL, y = NULL)

# Combining plots
p_ks_legumes <- wrap_plots(
    gmax_ks_distro +
        labs(title = "Glycine max") +
        theme(plot.title = element_text(face = "italic")),
    pvu_ks_distro +
        labs(title = "Phaseolus vulgaris") +
        theme(plot.title = element_text(face = "italic")),
    nrow = 1
)
```

```
p_ks_legumes
```

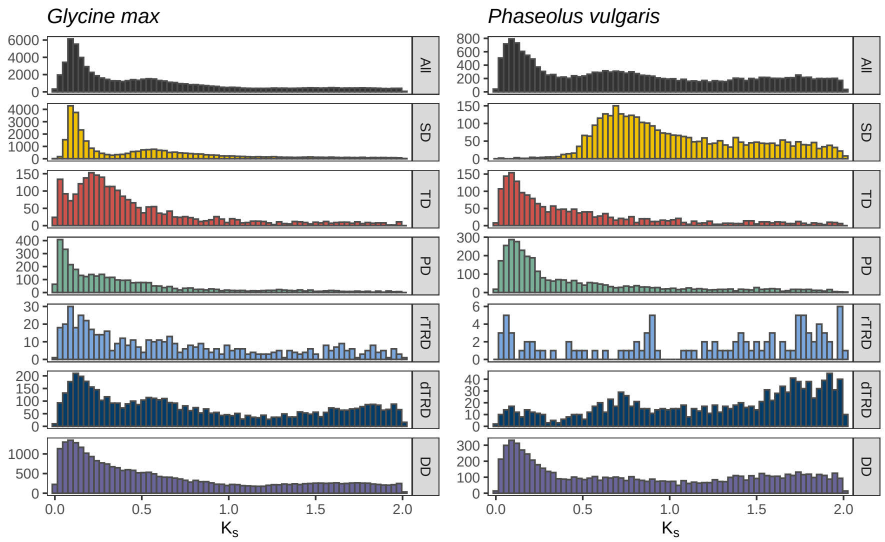

The plot shows the importance of visualizing Ks distributions by mode. When visualizing the whole-paranome distribution, detection of peaks is not trivial, and potential whole-genome duplication events might be masked. When we split the distribution by mode of duplication, we can more easily observe segmental duplicates that cluster together, providing strong evidence for whole-genome duplication events (2 events for *G. max*, and 1 events for *P. vulgaris*).

Next, we will plot the distributions of $K_a$, $K_s$, and $K_a/K_s$ values for selected plant species with phylogenetic context.

```
# Subset plant tree to get selected species only
tree_subset <- ape::keep.tip(tree_plants, names(plants_kaks))

# Clean names
names(plants_kaks) <- gsub("_", " ", str_to_title(names(plants_kaks)))
tree_subset$tip.label <- gsub("_", " ", str_to_title(tree_subset$tip.label))

# Plot tree
p_tree_selected <- ggtree(tree_subset, branch.length = "none") +
    geom_tiplab(fontface = "italic", size = 3)

# Reoder rates list based on tree topology
ord <- rev(ggtree::get_taxa_name(p_tree_selected))
rl <- plants_kaks[ord]

# Plot rates by species with tree on the left
p_rates_phylogeny <- wrap_plots(
    p_tree_selected + xlim(0, 10),
    plot_rates_by_species(rl, rate_column = "Ks", range = c(0, 2)) +
        theme(
            axis.text.y = element_blank(),
            axis.ticks.y = element_blank()
```

```
        ),
    plot_rates_by_species(
        rl, rate_column = "Ka", range = c(0, 2),
        fill = "mediumseagreen", color = "seagreen"
    ) +
        theme(
            axis.text.y = element_blank(),
            axis.ticks.y = element_blank()
        ),
    plot_rates_by_species(
        rl, rate_column = "Ka_Ks", range = c(0, 2),
        fill = "darkorange2", color = "darkorange3"
    ) +
        theme(
            axis.text.y = element_blank(),
            axis.ticks.y = element_blank()
        ),
    nrow = 1
) +
    plot_annotation(title = "Substitution rates in a phylogenetic context")
```

```
p_rates_phylogeny
```

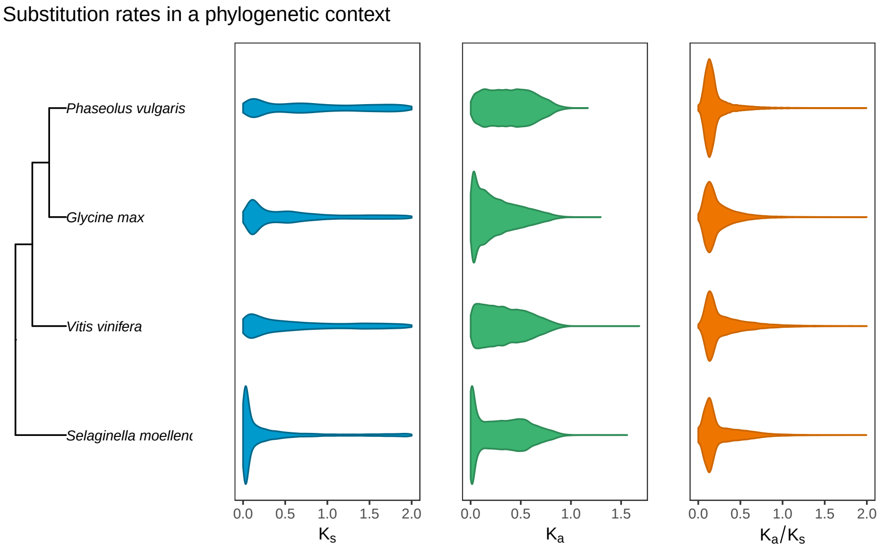

### Saving objects

Finally, let's save important objects created in this session for further use.

```
# Save plots for each instance
save(
    p_fungi, compress = "xz",
    file = here("products", "plots", "p_fungi.rda")
```

```
        )

        save(
            p_metazoa, compress = "xz",
            file = here("products", "plots", "p_metazoa.rda")
        )

        save(
            p_protists, compress = "xz",
            file = here("products", "plots", "p_protists.rda")
        )

        save(
            p_vertebrates, compress = "xz",
            file = here("products", "plots", "p_vertebrates.rda")
        )

        save(
            p_plants, compress = "xz",
            file = here("products", "plots", "p_plants.rda")
        )

        save(
            p_duplicates_all_ensembl, compress = "xz",
            file = here("products", "plots", "p_duplicates_all_ensembl.rda")
        )

        save(
            p_ks_legumes, compress = "xz",
            file = here("products", "plots", "p_ks_legumes.rda")
        )

        save(
            p_rates_phylogeny, compress = "xz",
            file = here("products", "plots", "p_rates_phylogeny.rda")
        )

        save(
            p_busco_association, compress = "xz",
            file = here("products", "plots", "p_busco_association.rda")
        )

        # Save tables
        save(
            sd_abundant_spp, compress = "xz",
            file = here("products", "result_files", "sd_abundant_spp.rda")
        )
```

### Session info

This document was created under the following conditions:

```
## - Session info ---------------------------------------------------------------
##  setting  value
##  version  R version 4.3.2 (2023-10-31)
##  os       Ubuntu 22.04.3 LTS
##  system   x86_64, linux-gnu
##  ui       X11
##  language (EN)
##  collate  en_US.UTF-8
##  ctype    en_US.UTF-8
##  tz       Europe/Brussels
##  date     2024-02-27
##  pandoc   3.1.1 @ /usr/lib/rstudio/resources/app/bin/quarto/bin/tools/ (via rmarkdown)
##
## - Packages ---------------------------------------------------------------------
##  package              * version     date (UTC) lib source
##  abind                  1.4-5       2016-07-21 [1] CRAN (R 4.3.2)
##  ade4                   1.7-22      2023-02-06 [1] CRAN (R 4.3.2)
##  AnnotationDbi          1.64.1      2023-11-03 [1] Bioconductor
##  ape                    5.7-1       2023-03-13 [1] CRAN (R 4.3.2)
##  aplot                  0.2.2       2023-10-06 [1] CRAN (R 4.3.2)
##  backports              1.4.1       2021-12-13 [1] CRAN (R 4.3.2)
##  Biobase                2.62.0      2023-10-24 [1] Bioconductor
##  BiocFileCache          2.10.1      2023-10-26 [1] Bioconductor
##  BiocGenerics           0.48.1      2023-11-01 [1] Bioconductor
##  BiocIO                 1.12.0      2023-10-24 [1] Bioconductor
##  BiocManager            1.30.22     2023-08-08 [1] CRAN (R 4.3.2)
##  BiocParallel           1.37.0      2024-01-19 [1] Github (Bioconductor/BiocParallel@79a1b2d)
##  BiocStyle            * 2.30.0      2023-10-24 [1] Bioconductor
##  biomaRt                2.58.2      2024-01-30 [1] Bioconductor 3.18 (R 4.3.2)
##  Biostrings             2.70.2      2024-01-28 [1] Bioconductor 3.18 (R 4.3.2)
##  bit                    4.0.5       2022-11-15 [1] CRAN (R 4.3.2)
##  bit64                  4.0.5       2020-08-30 [1] CRAN (R 4.3.2)
##  bitops                 1.0-7       2021-04-24 [1] CRAN (R 4.3.2)
##  blob                   1.2.4       2023-03-17 [1] CRAN (R 4.3.2)
##  bookdown               0.37        2023-12-01 [1] CRAN (R 4.3.2)
##  broom                  1.0.5       2023-06-09 [1] CRAN (R 4.3.2)
##  cachem                 1.0.8       2023-05-01 [1] CRAN (R 4.3.2)
##  car                    3.1-2       2023-03-30 [1] CRAN (R 4.3.2)
##  carData                3.0-5       2022-01-06 [1] CRAN (R 4.3.2)
##  cli                    3.6.2       2023-12-11 [1] CRAN (R 4.3.2)
##  coda                   0.19-4.1    2024-01-31 [1] CRAN (R 4.3.2)
##  codetools              0.2-19      2023-02-01 [4] CRAN (R 4.2.2)
##  colorspace             2.1-0       2023-01-23 [1] CRAN (R 4.3.2)
##  crayon                 1.5.2       2022-09-29 [1] CRAN (R 4.3.2)
##  curl                   5.2.0       2023-12-08 [1] CRAN (R 4.3.2)
##  DBI                    1.2.1       2024-01-12 [1] CRAN (R 4.3.2)
##  dbplyr                 2.4.0       2023-10-26 [1] CRAN (R 4.3.2)
```

```
##  DelayedArray           0.28.0     2023-10-24 [1] Bioconductor
##  digest                 0.6.34     2024-01-11 [1] CRAN (R 4.3.2)
##  doParallel             1.0.17     2022-02-07 [1] CRAN (R 4.3.2)
##  doubletrouble        * 1.3.4      2024-02-05 [1] Bioconductor
##  dplyr                * 1.1.4      2023-11-17 [1] CRAN (R 4.3.2)
##  evaluate               0.23       2023-11-01 [1] CRAN (R 4.3.2)
##  fansi                  1.0.6      2023-12-08 [1] CRAN (R 4.3.2)
##  farver                 2.1.1      2022-07-06 [1] CRAN (R 4.3.2)
##  fastmap                1.1.1      2023-02-24 [1] CRAN (R 4.3.2)
##  filelock               1.0.3      2023-12-11 [1] CRAN (R 4.3.2)
##  forcats              * 1.0.0      2023-01-29 [1] CRAN (R 4.3.2)
##  foreach                1.5.2      2022-02-02 [1] CRAN (R 4.3.2)
##  fs                     1.6.3      2023-07-20 [1] CRAN (R 4.3.2)
##  generics               0.1.3      2022-07-05 [1] CRAN (R 4.3.2)
##  GenomeInfoDb           1.38.6     2024-02-08 [1] Bioconductor 3.18 (R 4.3.2)
##  GenomeInfoDbData       1.2.11     2023-12-21 [1] Bioconductor
##  GenomicAlignments      1.38.2     2024-01-16 [1] Bioconductor 3.18 (R 4.3.2)
##  GenomicFeatures        1.54.3     2024-01-31 [1] Bioconductor 3.18 (R 4.3.2)
##  GenomicRanges          1.54.1     2023-10-29 [1] Bioconductor
##  ggfun                  0.1.4      2024-01-19 [1] CRAN (R 4.3.2)
##  ggnetwork              0.5.13     2024-02-14 [1] CRAN (R 4.3.2)
##  ggplot2              * 3.4.4      2023-10-12 [1] CRAN (R 4.3.2)
##  ggplotify              0.1.2      2023-08-09 [1] CRAN (R 4.3.2)
##  ggpubr                 0.6.0.999  2024-02-09 [1] Github (kassambara/ggpubr@6aeb4f7)
##  ggsignif               0.6.4      2022-10-13 [1] CRAN (R 4.3.2)
##  ggtree               * 3.10.0     2023-10-24 [1] Bioconductor
##  glue                   1.7.0      2024-01-09 [1] CRAN (R 4.3.2)
##  gridGraphics           0.5-1      2020-12-13 [1] CRAN (R 4.3.2)
##  gtable                 0.3.4      2023-08-21 [1] CRAN (R 4.3.2)
##  here                 * 1.0.1      2020-12-13 [1] CRAN (R 4.3.2)
##  hms                    1.1.3      2023-03-21 [1] CRAN (R 4.3.2)
##  htmltools              0.5.7      2023-11-03 [1] CRAN (R 4.3.2)
##  htmlwidgets            1.6.4      2023-12-06 [1] CRAN (R 4.3.2)
##  httr                   1.4.7      2023-08-15 [1] CRAN (R 4.3.2)
##  igraph                 2.0.1.1    2024-01-30 [1] CRAN (R 4.3.2)
##  intergraph             2.0-4      2024-02-01 [1] CRAN (R 4.3.2)
##  IRanges                2.36.0     2023-10-24 [1] Bioconductor
##  iterators              1.0.14     2022-02-05 [1] CRAN (R 4.3.2)
##  jsonlite               1.8.8      2023-12-04 [1] CRAN (R 4.3.2)
##  KEGGREST               1.42.0     2023-10-24 [1] Bioconductor
##  knitr                  1.45       2023-10-30 [1] CRAN (R 4.3.2)
##  labeling               0.4.3      2023-08-29 [1] CRAN (R 4.3.2)
##  lattice                0.22-5     2023-10-24 [4] CRAN (R 4.3.1)
##  lazyeval               0.2.2      2019-03-15 [1] CRAN (R 4.3.2)
##  lifecycle              1.0.4      2023-11-07 [1] CRAN (R 4.3.2)
##  lubridate            * 1.9.3      2023-09-27 [1] CRAN (R 4.3.2)
##  magrittr               2.0.3      2022-03-30 [1] CRAN (R 4.3.2)
##  MASS                   7.3-60     2023-05-04 [4] CRAN (R 4.3.1)
##  Matrix                 1.6-3      2023-11-14 [4] CRAN (R 4.3.2)
##  MatrixGenerics         1.14.0     2023-10-24 [1] Bioconductor
##  matrixStats            1.2.0      2023-12-11 [1] CRAN (R 4.3.2)
```

```
##  mclust                  6.0.1      2023-11-15 [1] CRAN (R 4.3.2)
##  memoise                 2.0.1      2021-11-26 [1] CRAN (R 4.3.2)
##  mgcv                    1.9-0      2023-07-11 [4] CRAN (R 4.3.1)
##  MSA2dist                1.6.0      2023-10-24 [1] Bioconductor
##  munsell                 0.5.0      2018-06-12 [1] CRAN (R 4.3.2)
##  network                 1.18.2     2023-12-05 [1] CRAN (R 4.3.2)
##  networkD3               0.4        2017-03-18 [1] CRAN (R 4.3.2)
##  nlme                    3.1-163    2023-08-09 [4] CRAN (R 4.3.1)
##  patchwork             * 1.2.0      2024-01-08 [1] CRAN (R 4.3.2)
##  pheatmap                1.0.12     2019-01-04 [1] CRAN (R 4.3.2)
##  pillar                  1.9.0      2023-03-22 [1] CRAN (R 4.3.2)
##  pkgconfig               2.0.3      2019-09-22 [1] CRAN (R 4.3.2)
##  png                     0.1-8      2022-11-29 [1] CRAN (R 4.3.2)
##  prettyunits             1.2.0      2023-09-24 [1] CRAN (R 4.3.2)
##  progress                1.2.3      2023-12-06 [1] CRAN (R 4.3.2)
##  purrr                 * 1.0.2      2023-08-10 [1] CRAN (R 4.3.2)
##  R6                      2.5.1      2021-08-19 [1] CRAN (R 4.3.2)
##  rappdirs                0.3.3      2021-01-31 [1] CRAN (R 4.3.2)
##  RColorBrewer            1.1-3      2022-04-03 [1] CRAN (R 4.3.2)
##  Rcpp                    1.0.12     2024-01-09 [1] CRAN (R 4.3.2)
##  RCurl                   1.98-1.14  2024-01-09 [1] CRAN (R 4.3.2)
##  readr                 * 2.1.5      2024-01-10 [1] CRAN (R 4.3.2)
##  restfulr                0.0.15     2022-06-16 [1] CRAN (R 4.3.2)
##  rjson                   0.2.21     2022-01-09 [1] CRAN (R 4.3.2)
##  rlang                   1.1.3      2024-01-10 [1] CRAN (R 4.3.2)
##  rmarkdown               2.25       2023-09-18 [1] CRAN (R 4.3.2)
##  rprojroot               2.0.4      2023-11-05 [1] CRAN (R 4.3.2)
##  Rsamtools               2.18.0     2023-10-24 [1] Bioconductor
##  RSQLite                 2.3.5      2024-01-21 [1] CRAN (R 4.3.2)
##  rstatix                 0.7.2      2023-02-01 [1] CRAN (R 4.3.2)
##  rstudioapi              0.15.0     2023-07-07 [1] CRAN (R 4.3.2)
##  rtracklayer             1.62.0     2023-10-24 [1] Bioconductor
##  S4Arrays                1.2.0      2023-10-24 [1] Bioconductor
##  S4Vectors               0.40.2     2023-11-23 [1] Bioconductor 3.18 (R 4.3.2)
##  scales                  1.3.0      2023-11-28 [1] CRAN (R 4.3.2)
##  seqinr                  4.2-36     2023-12-08 [1] CRAN (R 4.3.2)
##  sessioninfo             1.2.2      2021-12-06 [1] CRAN (R 4.3.2)
##  SparseArray             1.2.4      2024-02-11 [1] Bioconductor 3.18 (R 4.3.2)
##  statnet.common          4.9.0      2023-05-24 [1] CRAN (R 4.3.2)
##  stringi                 1.8.3      2023-12-11 [1] CRAN (R 4.3.2)
##  stringr               * 1.5.1      2023-11-14 [1] CRAN (R 4.3.2)
##  SummarizedExperiment    1.32.0     2023-10-24 [1] Bioconductor
##  syntenet                1.4.0      2023-10-24 [1] Bioconductor
##  tibble                * 3.2.1      2023-03-20 [1] CRAN (R 4.3.2)
##  tidyr                 * 1.3.1      2024-01-24 [1] CRAN (R 4.3.2)
##  tidyselect              1.2.0      2022-10-10 [1] CRAN (R 4.3.2)
##  tidytree                0.4.6      2023-12-12 [1] CRAN (R 4.3.2)
##  tidyverse             * 2.0.0      2023-02-22 [1] CRAN (R 4.3.2)
##  timechange              0.3.0      2024-01-18 [1] CRAN (R 4.3.2)
##  treeio                  1.26.0     2023-10-24 [1] Bioconductor
##  tzdb                    0.4.0      2023-05-12 [1] CRAN (R 4.3.2)
```

```
##  utf8                   1.2.4      2023-10-22 [1] CRAN (R 4.3.2)
##  vctrs                  0.6.5      2023-12-01 [1] CRAN (R 4.3.2)
##  withr                  3.0.0      2024-01-16 [1] CRAN (R 4.3.2)
##  xfun                   0.42       2024-02-08 [1] CRAN (R 4.3.2)
##  XML                    3.99-0.16.1 2024-01-22 [1] CRAN (R 4.3.2)
##  xml2                   1.3.6      2023-12-04 [1] CRAN (R 4.3.2)
##  XVector                0.42.0     2023-10-24 [1] Bioconductor
##  yaml                   2.3.8      2023-12-11 [1] CRAN (R 4.3.2)
##  yulab.utils            0.1.4      2024-01-28 [1] CRAN (R 4.3.2)
##  zlibbioc               1.48.0     2023-10-24 [1] Bioconductor
##
##  [1] /home/faalm/R/x86_64-pc-linux-gnu-library/4.3
##  [2] /usr/local/lib/R/site-library
##  [3] /usr/lib/R/site-library
##  [4] /usr/lib/R/library
##
## ------------------------------------------------------------------------------
```

### Supplementary Text S5: Runtime benchmark

***Fabricio Almeida-Silva*** [1] ***and Yves Van de Peer*** [1]

[1]VIB-UGent Center for Plant Systems Biology, Ghent University, Ghent, Belgium

**27 February 2024**

### 1 Introduction

Here, we will perform a runtime benchmark for functions related to duplicate classification and substitution rates calculation using model organisms.

To start, let's load the required data and packages.

```
set.seed(123) # for reproducibility

# Load required packages
library(doubletrouble)
library(here)
## here() starts at /home/faalm/Dropbox/package_benchmarks/doubletrouble_paper
library(tidyverse)
## -- Attaching core tidyverse packages ------------------------ tidyverse 2.0.0 --
## v dplyr     1.1.4     v readr     2.1.5
## v forcats   1.0.0     v stringr   1.5.1
## v ggplot2   3.4.4     v tibble    3.2.1
## v lubridate 1.9.3     v tidyr     1.3.1
## v purrr     1.0.2
## -- Conflicts ------------------------------------------ tidyverse_conflicts() --
## x dplyr::filter() masks stats::filter()
## x dplyr::lag()    masks stats::lag()
## i Use the conflicted package (<http://conflicted.r-lib.org/>) to force all conflicts to become errors
library(patchwork)

source(here("code", "utils.R"))

# Load sample metadata for Ensembl instances
load(here("products", "result_files", "metadata_all.rda"))
```

### 2 Benchmark 1: `classify_gene_pairs()`

Here, we will benchmark the performance of `classify_gene_pairs()` with model organisms.

First, let's get the genome and annotation data.

```
# Create a data frame with names of model species and their Ensembl instances
model_species <- data.frame(
    species = c(
        "arabidopsis_thaliana", "caenorhabditis_elegans",
        "homo_sapiens", "saccharomyces_cerevisiae",
        "drosophila_melanogaster", "danio_rerio"
    ),
    instance = c(
        "plants", "metazoa", "ensembl", "fungi", "metazoa", "ensembl"
    )
)

# For each organism, download data, and identify and classify duplicates
model_duplicates <- lapply(seq_len(nrow(model_species)), function(x) {
```

```r
        species <- model_species$species[x]
        instance <- model_species$instance[x]

        # Get annotation
        annot <- get_annotation(model_species[x, ], instance)

        # Get proteome and keep only primary transcripts
        seq <- get_proteomes(model_species[x, ], instance)
        seq <- filter_sequences(seq, annot)

        # Process data
        pdata <- syntenet::process_input(seq, annot, filter_annotation = TRUE)

        # Perform DIAMOND search
        outdir <- file.path(tempdir(), paste0(species, "_intra"))
        diamond <- syntenet::run_diamond(
            seq = pdata$seq,
            compare = "intraspecies",
            outdir = outdir,
            threads = 4,
            ... = "--sensitive"
        )

        fs::dir_delete(outdir)

        # Classify duplicates - standard mode
        start <- Sys.time()
        duplicate_pairs <- classify_gene_pairs(
                blast_list = diamond,
                annotation = pdata$annotation,
                scheme = "standard"
            )[[1]]
        end <- Sys.time()
        runtime <- end - start

        return(runtime)
    })
    names(model_duplicates) <- gsub("_", " ", str_to_title(model_species$species))

    # Summarize results in a table
    benchmark_classification <- data.frame(
        species = names(model_duplicates),
        time_seconds = as.numeric(unlist(model_duplicates))
    )

    # Save results
    save(
        benchmark_classification, compress = "xz",
        file = here("products", "result_files", "benchmark_classification.rda")
    )
```

### 3 Benchmark 2: `pairs2kaks()`

Next, we will benchmark the performance of `pairs2kaks()` for duplicate pairs in the *Saccharomyces cerevisiae* genome. We will do it using a single thread, and using parallelization (with 4 and 8 threads).

First of all, let's get the required data for `pairs2kaks()`.

```
# Load duplicate pairs for S. cerevisiae
load(here("products", "result_files", "fungi_duplicates.rda"))
scerevisiae_pairs <- fungi_duplicates["saccharomyces_cerevisiae"]

# Get CDS for S. cerevisiae
scerevisiae_cds <- get_cds_ensembl("saccharomyces_cerevisiae", "fungi")
```

Now, we can do the benchmark.

```
# Parallel back-end: SerialParam (1 thread)
start <- Sys.time()
kaks <- pairs2kaks(
    scerevisiae_pairs,
    scerevisiae_cds,
    bp_param = BiocParallel::SerialParam()
)
end <- Sys.time()
runtime_serial <- end - start

# Parallel back-end: SnowParam, 4 threads
start <- Sys.time()
kaks <- pairs2kaks(
    scerevisiae_pairs,
    scerevisiae_cds,
    bp_param = BiocParallel::SnowParam(workers = 4)
)
end <- Sys.time()
runtime_snow4 <- end - start

# Parallel back-end: SnowParam, 8 threads
start <- Sys.time()
kaks <- pairs2kaks(
    scerevisiae_pairs,
    scerevisiae_cds,
    bp_param = BiocParallel::SnowParam(workers = 8)
)
end <- Sys.time()
runtime_snow8 <- end - start

# Summarize results in a table
benchmark_kaks <- data.frame(
    `Back-end` = c("Serial", "Snow, 4 threads", "Snow, 8 threads"),
    Time_minutes = as.numeric(c(runtime_serial, runtime_snow4, runtime_snow8))
) |>
```

```
    dplyr::mutate(
        Pairs_per_minute = nrow(scerevisiae_pairs[[1]]) / Time_minutes,
        Pairs_per_second = nrow(scerevisiae_pairs[[1]]) / (Time_minutes * 60)
    )

save(
    benchmark_kaks, compress = "xz",
    file = here("products", "result_files", "benchmark_kaks.rda")
)
```

### Session info

This document was created under the following conditions:

```
## - Session info ---------------------------------------------------------------
##  setting  value
##  version  R version 4.3.2 (2023-10-31)
##  os       Ubuntu 22.04.3 LTS
##  system   x86_64, linux-gnu
##  ui       X11
##  language (EN)
##  collate  en_US.UTF-8
##  ctype    en_US.UTF-8
##  tz       Europe/Brussels
##  date     2024-02-27
##  pandoc   3.1.1 @ /usr/lib/rstudio/resources/app/bin/quarto/bin/tools/ (via rmarkdown)
##
## - Packages -------------------------------------------------------------------
##  package              * version     date (UTC) lib source
##  abind                  1.4-5       2016-07-21 [1] CRAN (R 4.3.2)
##  ade4                   1.7-22      2023-02-06 [1] CRAN (R 4.3.2)
##  AnnotationDbi          1.64.1      2023-11-03 [1] Bioconductor
##  ape                    5.7-1       2023-03-13 [1] CRAN (R 4.3.2)
##  Biobase                2.62.0      2023-10-24 [1] Bioconductor
##  BiocFileCache          2.10.1      2023-10-26 [1] Bioconductor
##  BiocGenerics           0.48.1      2023-11-01 [1] Bioconductor
##  BiocIO                 1.12.0      2023-10-24 [1] Bioconductor
##  BiocManager            1.30.22     2023-08-08 [1] CRAN (R 4.3.2)
##  BiocParallel           1.37.0      2024-01-19 [1] Github (Bioconductor/BiocParallel@79a1b2d)
##  BiocStyle            * 2.30.0      2023-10-24 [1] Bioconductor
##  biomaRt                2.58.2      2024-01-30 [1] Bioconductor 3.18 (R 4.3.2)
##  Biostrings             2.70.2      2024-01-28 [1] Bioconductor 3.18 (R 4.3.2)
##  bit                    4.0.5       2022-11-15 [1] CRAN (R 4.3.2)
##  bit64                  4.0.5       2020-08-30 [1] CRAN (R 4.3.2)
##  bitops                 1.0-7       2021-04-24 [1] CRAN (R 4.3.2)
##  blob                   1.2.4       2023-03-17 [1] CRAN (R 4.3.2)
##  bookdown               0.37        2023-12-01 [1] CRAN (R 4.3.2)
##  cachem                 1.0.8       2023-05-01 [1] CRAN (R 4.3.2)
##  cli                    3.6.2       2023-12-11 [1] CRAN (R 4.3.2)
```

```
##  coda                  0.19-4.1   2024-01-31 [1] CRAN (R 4.3.2)
##  codetools             0.2-19     2023-02-01 [4] CRAN (R 4.2.2)
##  colorspace            2.1-0      2023-01-23 [1] CRAN (R 4.3.2)
##  crayon                1.5.2      2022-09-29 [1] CRAN (R 4.3.2)
##  curl                  5.2.0      2023-12-08 [1] CRAN (R 4.3.2)
##  DBI                   1.2.1      2024-01-12 [1] CRAN (R 4.3.2)
##  dbplyr                2.4.0      2023-10-26 [1] CRAN (R 4.3.2)
##  DelayedArray          0.28.0     2023-10-24 [1] Bioconductor
##  digest                0.6.34     2024-01-11 [1] CRAN (R 4.3.2)
##  doParallel            1.0.17     2022-02-07 [1] CRAN (R 4.3.2)
##  doubletrouble       * 1.3.4      2024-02-05 [1] Bioconductor
##  dplyr               * 1.1.4      2023-11-17 [1] CRAN (R 4.3.2)
##  evaluate              0.23       2023-11-01 [1] CRAN (R 4.3.2)
##  fansi                 1.0.6      2023-12-08 [1] CRAN (R 4.3.2)
##  fastmap               1.1.1      2023-02-24 [1] CRAN (R 4.3.2)
##  filelock              1.0.3      2023-12-11 [1] CRAN (R 4.3.2)
##  forcats             * 1.0.0      2023-01-29 [1] CRAN (R 4.3.2)
##  foreach               1.5.2      2022-02-02 [1] CRAN (R 4.3.2)
##  generics              0.1.3      2022-07-05 [1] CRAN (R 4.3.2)
##  GenomeInfoDb          1.38.6     2024-02-08 [1] Bioconductor 3.18 (R 4.3.2)
##  GenomeInfoDbData      1.2.11     2023-12-21 [1] Bioconductor
##  GenomicAlignments     1.38.2     2024-01-16 [1] Bioconductor 3.18 (R 4.3.2)
##  GenomicFeatures       1.54.3     2024-01-31 [1] Bioconductor 3.18 (R 4.3.2)
##  GenomicRanges         1.54.1     2023-10-29 [1] Bioconductor
##  ggnetwork             0.5.13     2024-02-14 [1] CRAN (R 4.3.2)
##  ggplot2             * 3.4.4      2023-10-12 [1] CRAN (R 4.3.2)
##  glue                  1.7.0      2024-01-09 [1] CRAN (R 4.3.2)
##  gtable                0.3.4      2023-08-21 [1] CRAN (R 4.3.2)
##  here                * 1.0.1      2020-12-13 [1] CRAN (R 4.3.2)
##  hms                   1.1.3      2023-03-21 [1] CRAN (R 4.3.2)
##  htmltools             0.5.7      2023-11-03 [1] CRAN (R 4.3.2)
##  htmlwidgets           1.6.4      2023-12-06 [1] CRAN (R 4.3.2)
##  httr                  1.4.7      2023-08-15 [1] CRAN (R 4.3.2)
##  igraph                2.0.1.1    2024-01-30 [1] CRAN (R 4.3.2)
##  intergraph            2.0-4      2024-02-01 [1] CRAN (R 4.3.2)
##  IRanges               2.36.0     2023-10-24 [1] Bioconductor
##  iterators             1.0.14     2022-02-05 [1] CRAN (R 4.3.2)
##  KEGGREST              1.42.0     2023-10-24 [1] Bioconductor
##  knitr                 1.45       2023-10-30 [1] CRAN (R 4.3.2)
##  lattice               0.22-5     2023-10-24 [4] CRAN (R 4.3.1)
##  lifecycle             1.0.4      2023-11-07 [1] CRAN (R 4.3.2)
##  lubridate           * 1.9.3      2023-09-27 [1] CRAN (R 4.3.2)
##  magrittr              2.0.3      2022-03-30 [1] CRAN (R 4.3.2)
##  MASS                  7.3-60     2023-05-04 [4] CRAN (R 4.3.1)
##  Matrix                1.6-3      2023-11-14 [4] CRAN (R 4.3.2)
##  MatrixGenerics        1.14.0     2023-10-24 [1] Bioconductor
##  matrixStats           1.2.0      2023-12-11 [1] CRAN (R 4.3.2)
##  mclust                6.0.1      2023-11-15 [1] CRAN (R 4.3.2)
##  memoise               2.0.1      2021-11-26 [1] CRAN (R 4.3.2)
##  MSA2dist              1.6.0      2023-10-24 [1] Bioconductor
##  munsell               0.5.0      2018-06-12 [1] CRAN (R 4.3.2)
```

```
##  network                  1.18.2      2023-12-05 [1] CRAN (R 4.3.2)
##  networkD3                0.4         2017-03-18 [1] CRAN (R 4.3.2)
##  nlme                     3.1-163     2023-08-09 [4] CRAN (R 4.3.1)
##  patchwork              * 1.2.0       2024-01-08 [1] CRAN (R 4.3.2)
##  pheatmap                 1.0.12      2019-01-04 [1] CRAN (R 4.3.2)
##  pillar                   1.9.0       2023-03-22 [1] CRAN (R 4.3.2)
##  pkgconfig                2.0.3       2019-09-22 [1] CRAN (R 4.3.2)
##  png                      0.1-8       2022-11-29 [1] CRAN (R 4.3.2)
##  prettyunits              1.2.0       2023-09-24 [1] CRAN (R 4.3.2)
##  progress                 1.2.3       2023-12-06 [1] CRAN (R 4.3.2)
##  purrr                  * 1.0.2       2023-08-10 [1] CRAN (R 4.3.2)
##  R6                       2.5.1       2021-08-19 [1] CRAN (R 4.3.2)
##  rappdirs                 0.3.3       2021-01-31 [1] CRAN (R 4.3.2)
##  RColorBrewer             1.1-3       2022-04-03 [1] CRAN (R 4.3.2)
##  Rcpp                     1.0.12      2024-01-09 [1] CRAN (R 4.3.2)
##  RCurl                    1.98-1.14   2024-01-09 [1] CRAN (R 4.3.2)
##  readr                  * 2.1.5       2024-01-10 [1] CRAN (R 4.3.2)
##  restfulr                 0.0.15      2022-06-16 [1] CRAN (R 4.3.2)
##  rjson                    0.2.21      2022-01-09 [1] CRAN (R 4.3.2)
##  rlang                    1.1.3       2024-01-10 [1] CRAN (R 4.3.2)
##  rmarkdown                2.25        2023-09-18 [1] CRAN (R 4.3.2)
##  rprojroot                2.0.4       2023-11-05 [1] CRAN (R 4.3.2)
##  Rsamtools                2.18.0      2023-10-24 [1] Bioconductor
##  RSQLite                  2.3.5       2024-01-21 [1] CRAN (R 4.3.2)
##  rstudioapi               0.15.0      2023-07-07 [1] CRAN (R 4.3.2)
##  rtracklayer              1.62.0      2023-10-24 [1] Bioconductor
##  S4Arrays                 1.2.0       2023-10-24 [1] Bioconductor
##  S4Vectors                0.40.2      2023-11-23 [1] Bioconductor 3.18 (R 4.3.2)
##  scales                   1.3.0       2023-11-28 [1] CRAN (R 4.3.2)
##  seqinr                   4.2-36      2023-12-08 [1] CRAN (R 4.3.2)
##  sessioninfo              1.2.2       2021-12-06 [1] CRAN (R 4.3.2)
##  SparseArray              1.2.4       2024-02-11 [1] Bioconductor 3.18 (R 4.3.2)
##  statnet.common           4.9.0       2023-05-24 [1] CRAN (R 4.3.2)
##  stringi                  1.8.3       2023-12-11 [1] CRAN (R 4.3.2)
##  stringr                * 1.5.1       2023-11-14 [1] CRAN (R 4.3.2)
##  SummarizedExperiment     1.32.0      2023-10-24 [1] Bioconductor
##  syntenet                 1.4.0       2023-10-24 [1] Bioconductor
##  tibble                 * 3.2.1       2023-03-20 [1] CRAN (R 4.3.2)
##  tidyr                  * 1.3.1       2024-01-24 [1] CRAN (R 4.3.2)
##  tidyselect               1.2.0       2022-10-10 [1] CRAN (R 4.3.2)
##  tidyverse              * 2.0.0       2023-02-22 [1] CRAN (R 4.3.2)
##  timechange               0.3.0       2024-01-18 [1] CRAN (R 4.3.2)
##  tzdb                     0.4.0       2023-05-12 [1] CRAN (R 4.3.2)
##  utf8                     1.2.4       2023-10-22 [1] CRAN (R 4.3.2)
##  vctrs                    0.6.5       2023-12-01 [1] CRAN (R 4.3.2)
##  withr                    3.0.0       2024-01-16 [1] CRAN (R 4.3.2)
##  xfun                     0.42        2024-02-08 [1] CRAN (R 4.3.2)
##  XML                      3.99-0.16.1 2024-01-22 [1] CRAN (R 4.3.2)
##  xml2                     1.3.6       2023-12-04 [1] CRAN (R 4.3.2)
##  XVector                  0.42.0      2023-10-24 [1] Bioconductor
##  yaml                     2.3.8       2023-12-11 [1] CRAN (R 4.3.2)
```

```
##  zlibbioc               1.48.0      2023-10-24 [1] Bioconductor
##
##  [1] /home/faalm/R/x86_64-pc-linux-gnu-library/4.3
##  [2] /usr/local/lib/R/site-library
##  [3] /usr/lib/R/site-library
##  [4] /usr/lib/R/library
##
## ------------------------------------------------------------------------------
```
